# Genetic diversity of the Atacama Desert shrub *Huidobria chilensis* in the context of geography and climate

**DOI:** 10.1101/2023.09.20.558398

**Authors:** K. Bechir Ferchichi, T. Böhnert, B. Ritter, D. Harpke, A. Stoll, P. Morales, S. Fiedler, F. Mu, J. Bechteler, C. Münker, M.A. Koch, T. Wiehe, D. Quandt

**Affiliations:** Bonn Institute of Organismic Biodiversity (BIOB), University of Bonn, Meckenheimer Allee 170, 53115 Bonn, Germany; Crop Bioinformatics, University of Bonn, Bonn, 53115, Germany; Institute for Geology & Mineralogy, University of Cologne, Germany Zülpicher Str. 49b 50674 Cologne, Germany; Leibniz Institute of Plant Genetics and Crop Plant Research, Corrensstraße 3 06466 Gatersleben, Germany; University of La Serena -Andrés Bello Campus, Centro de Estudios Avanzados en Zonas Áridas (CEAZA), Raúl Bitrán 1305, La Serena Región de Coquimbo, Chile; GEOMAR Helmholtz Centre for Ocean Research Kiel, Düsternbrooker Weg 20 24105 Kiel Germany; University of Kaiserslautern-Landau (PRTU), Institute for Environmental Sciences, iES Landau, Fortstraße 7, 76829 Landau, Germany; University of Heidelberg, Centre for Organismal Studies (COS), Im Neuenheimer Feld 345, 69120 Heidelberg, Germany; University of Cologne, Institute for Genetics, Zülpicher Str. 47a, 50674 Cologne, Germany

**Keywords:** Speciation, Population Structure, Differentiation, Isolation by distance, Correlation

## Abstract

Survival in hyperarid deserts is a major challenge for plant life, requiring the development of evolutionary strategies. The Atacama Desert presents harsh conditions such as limited rainfall, crusted soils, high soil salinity, high altitude, and intense solar radiation. These conditions, together with paleoclimatic variability since the past millions of years, have influenced the genetic structure and connectivity of plant populations, resulting in a diverse flora with high endemism. However, the diversification of most lineages appears to be relatively recent, in contrast to proposed age of the Atacama Desert and the onset, evolution and expansion of hyperarid conditions since the Late Oligocene and Early Miocene. A prominent exception is the Atacama paleoendemic *Huidobria chilensis* (Loasaceae), which is thought to be adapted to such conditions since the Eocene. Still, the environmental limits and thresholds for life in the Atacama remain poorly understood. To investigate the genetic structure in relation to the history of the Atacama Desert, we studied 186 individuals from 11 populations using genotyping-by-sequencing (GBS). Genome-wide single nucleotide polymorphisms (SNPs) were analyzed for population structure and genetic diversity. We identified three genetic clusters corresponding to geographic regions: the coastal region south of Tocopilla, the Coastal Cordillera around Chañaral, and the Copiapó watershed in the south. These clusters as well as genetic diversity were analyzed alongside rainfall, altitude, and landscape data. Although the genetic data generally supports isolation by distance as a major factor for genetic variation between populations, the study also reveals the influence of the topography on the distribution of *H. chilensis* and highlights the role of hydrologically connected watersheds and rivers in plant migration and colonization. This shapes the species’ evolutionary trajectory and genetic diversity. Understanding these patterns provides insights into the adaptation and survival strategies of plants in extreme desert environments such as the Atacama.

## 1. Introduction

Ancient deserts, with their extreme and persistent environmental conditions, offer unparalleled opportunities to test theories of landscape and biological evolution. The low complexity and high aridity of these deserts provide a unique setting to study plant adaptation and survival strategies, shedding light on the fundamental mechanisms of adaptation and the limits of life on Earth (Stebbins, 1952). The Atacama Desert, with its prolonged hyperaridity (Dunai et al., 2005, Ritter et al. 2019b, Evenstar et al. 2017), sparse vegetation (Rundel et al., 1991) and complex landscape dynamics, stands out as an exceptional study site to investigate the interplay between evolutionary processes, landscape dynamics and plant responses (Dunai et al., 2020).

The Atacama Desert extends over 1000 km along the western coast of South America, from southern Peru (-18°S) to central Chile (-30°S), in a narrow strip between the Pacific Ocean in the west and the Andean Cordillera in the east (Gómez-Silva and Batista-García, 2022). The desert is influenced by two opposing meteorological systems, which cause low rainfall at different times of the year along the coast (from the west) and the Andes (from the east) (Houston et al. 2006). Cold upwelling waters along the coast cause the formation of a temperature inversion layer that traps moisture migration below about 1,000 to 1,200 m (Cereceda et al., 2008). This moisture creates coastal fog, so called ‘camanchaca’ (Cereceda et al. 2008), which creates local and rather low amounts of precipitation along the coast (Cereceda et al., 2008). Rare precipitation events, caused by cut-off low system can cause additional surface runoff in this region (Reyers et al. 2019, Reyers et al. 2020). Additional abiotic factors, such as crusted soils, high soil salinity, a strong altitudinal gradient, as well as high levels of UV radiation, pose serious challenges to plant survival in the Atacama Desert (Cordero et al., 2016; Sitzia et al., 2019).

However, these current climatic conditions have not been stable since the suggested Oligocene-Miocene onset of hyperaridity in parts of the desert and subsequent expansion of these conditions throughout the Atacama Desert (Dunai et al., 2005, Ritter et al. 2019, Evenstar et al. 2017), predominant hyperarid conditions exhibit also variability which reflect global trends of the paleoclimate throughout the Miocene (Zachos et al., 2001; Edwards et al., 2010, Ritter et al. 2018). Several palaeo-climatic models (Houston et al. 2006), as well as paleoclimate records (e.g. Ritter et al. 2019, Diederich et al. 2020, Medialdea et al. 2020, Jordan et al. 2014, Jordan et al. 2022 Ritter et al. 2022) have shown that during the Pleistocene and Pliocene, pluvial phases repeatedly interrupted the hyper-arid conditions. It has been repeatedly shown that this Pleistocene-Pliocene climate variability has shaped most of the plant species diversity (Luebert et al., 2008; Dillon et al., 2009; Böhnert et al., 2019) as well as biogeographic processes (Luebert et al., 2011; Böhnert et al., 2022).

Recent climate variability (e.g. Ritter et al. 2019, Diederich et al. 2020) has also been shown to have a strong impact on speciation events (Merklinger et al., 2021; Möbus et al., 2021) and population dynamics (Ossa et al., 2013; Merklinger et al., 2020). However, there are only a few population-level genomic studies for plants available for the Atacama Desert, all of which have a Quaternary or at least recent evolutionary history. In turn, no study has ever investigated whether the ancient climatic history has left a genomic imprint on palae-endemic lineages.

Few lineages can be considered palaeoendemic, however, a prominent example is the genus *Huidobria* Gay (Loasaceae), with two only distantly related species, *Huidobria fruticosa* Phil. and *Huidobria chilensis* Gay (Grau 1997; Acuña et al. 2019). Both species are shrubs with a distinctive growth habit and typical Loasaceae flowers (Weigend et al., 2004). *H. fruticosa* is more widespread, with distinct populations along the northern and southern coasts and in the Andes (Merklinger at al., submitted), while *H. chilensis* is found exclusively along the southern coast until Copiapoa (Fig. 1), where populations extend into the Andes (Grau 1997). Distributional overlap is restricted and so far, reports of joint occurrence is lacking. The species of focus in this study, *H. chilensis*, is smaller than its congener, with thinner, almost brittle leaves with a thick cuticle layer to prevent water loss and protect against UV radiation, presumably an adaptation to cope with desert conditions (Weigend et al., 2004). Like *H. fruticosa*, *H. chilensis* plants (Fig. 2) are almost exclusively found in catchment or drainage areas, but are generally absent from hillsides without access to groundwater, raising the question of the weighting of different abiotic factors responsible for its current distribution and genetic structure, as well as its abiotic delineation from *H. fruticosa*.

**Fig. 1.**
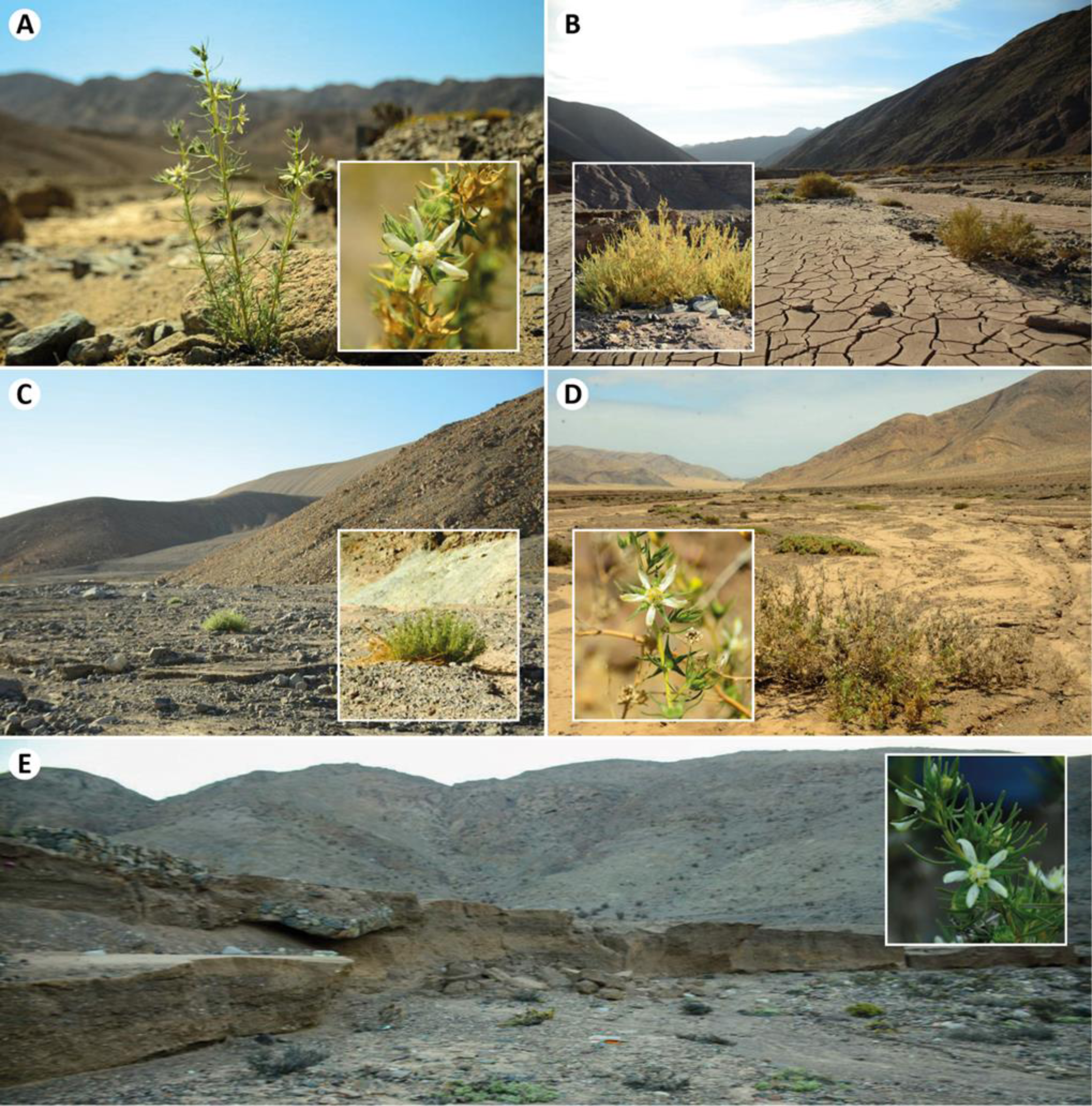
Captured habitat of *Huidobria chilensis* in five different sampling sites (Images from A. Stoll & P. Morales). A) Flowering H. *chilensis* in the southernmost location on the road C-386 to Copiapó (800). B) Growing *H. chilensis* on the road from Copiapó to the San Francisco International pass (801). C) One individual from the population 803 growing near to Quebrada (803). D) Flowering individuals of *H. chilensis* near to Cinfuncho on the road B-900 (808). E) Bed showing individuals distributed on the road 1 to Taltal (809).

**Fig. 2.**
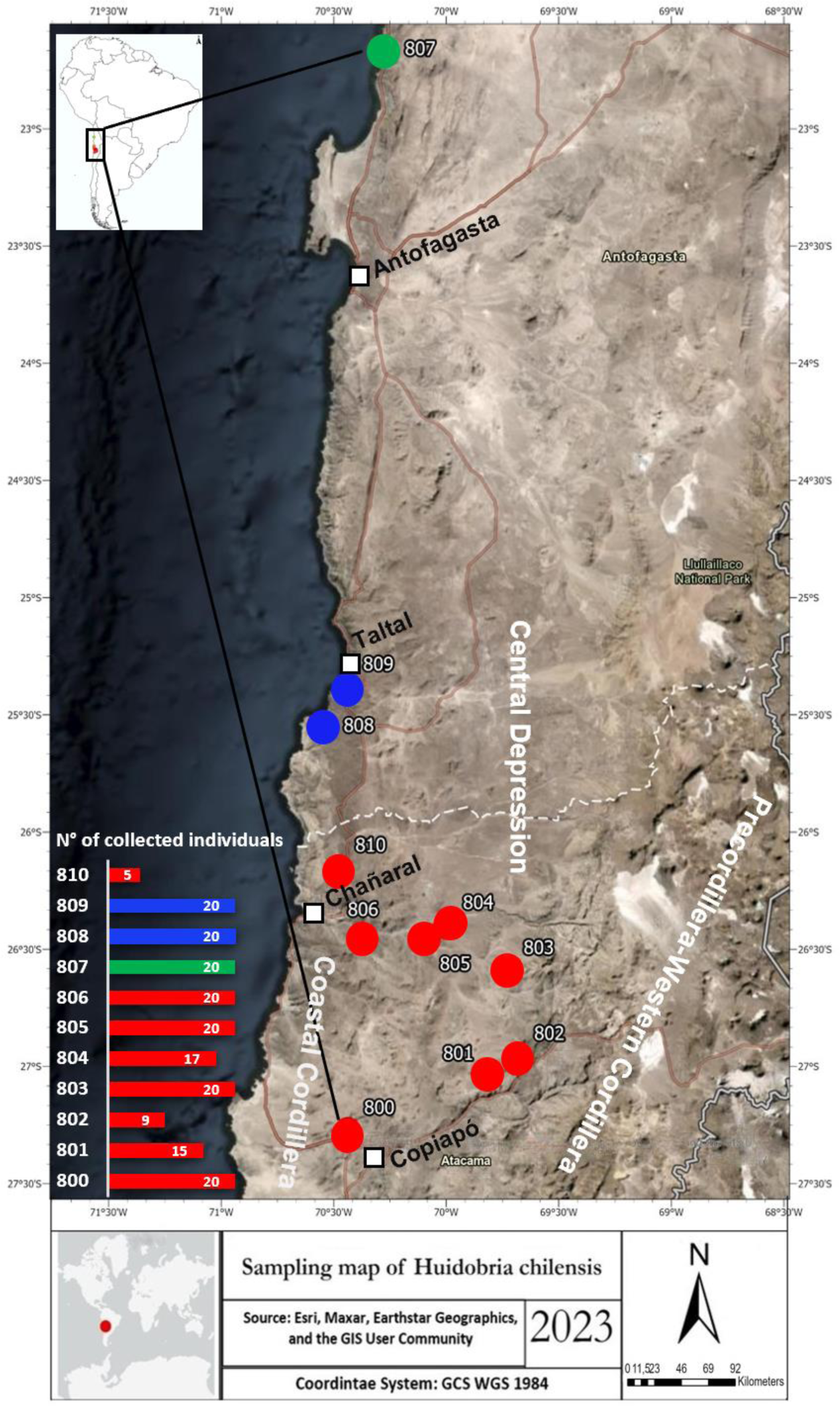
Overview map (based on Esri, HERE, Garmin, FAO, NOAA, USGS, © OpenStreetMap contributors, and the GIS User Community) generated by ArcGIS Pro v.3.1.0 and Adobe illustrator 2023 software of the study area. Sample collecting site are indicated in different colors according to their regions: Green (807): Coastal Cliff near Tocopilla; Blue (808 and 809): Coastal Cliff and Coastal Cordillera north of Chañaral; Red (800 to 806 and 810): Copiapó Watershed. All 11 locations are marked and presented by their ID numbers (800, 801, 802, 803, 804, 805, 806, 807, 808, 809, 810). The bar plot shows the number of individuals sampled per population.

Therefore, high-quality sequencing data were generated using modified genotyping-by-sequencing (GBS; Elshire et al., 2011) with two restriction enzymes (Peterson et al., 2012) to understand whether the long-term hyperarid conditions as well as the geological history of the Atacama Desert have left an imprint on the population structure. Such sequencing approaches belong to a family of closely related methods, often referred to as restriction site-associated DNA sequencing (RADseq), which are now widely used for population genomic studies (e.g., Andrews et al., 2016). In the present study, we seek to understand the influence of landscape and environmental features on the genetic diversity, population structure and current distribution of *H. chilensis*, in order to elucidate its evolutionary history. We postulated that variation in environmental factors may have acted as a driver of local adaptation, impeding gene flow between populations of the species. Therefore, we sought to investigate (i) the degree of genetic similarity between populations following their separation, (ii) the role of geographic isolation and environmental factors in genetically sub-structuring populations, and (iii) the ultimate cause of genetic differentiation between *H. chilensis* individuals in the regions studied.

## 2. Material and Methods

### 2.1. Field work and sampling

A total of 186 leaf samples of *Huidobria chilensis* were collected from 11 localities in the Atacama Desert of northern Chile during several field campaigns (Fig. 1; supplementary Table S6). In general, 25 individuals per population were sampled, except for sampling site 801 with 15, 802 with 9, 804 with 17 and 810 with only 5 individuals, respectively. The sampled leaves were directly silica dried. Vouchers for each population are available at the herbarium BONN (University of Bonn, Germany) and ULS (University of La Serena, Chile).

### 2.2. Plant DNA extraction

DNA isolation was performed from silica-dried leaf material using the Macherey Nagel Nucleo Mag 96 protocol following the manufacture’s protocol. The samples went through homogenization using lysis buffer before being loaded into 96 Square Well Blocks. The Thermo Fisher Scientific KingFisher Flex benchtop system was used to bind DNA to NucleoMag C-beads and eluted at 55 °C into 150 μl MC6 elution buffer. Lonza GelStar Nucleic Acid Gel Stain (100×) was used to assess the quality and quantity of the isolated products, while genomic DNA assessment was achieved by using a 20ng linear, double-stranded Lambda DNA probe (New England Biolabs, N3011S). Numerical values were assigned to the agarose-gel readings by measuring selected individuals using the Qubit 2.0 Fluorometer.

### 2.3. Genotyping-by-sequencing

200 ng of the genomic DNA was digested with *Pst*I-HF and *Msp*I restriction enzymes from New England Biolabs, USA. Adapters with unique barcodes were ligated to the resulting fragments using T4-ligase from the same supplier. Size-selection targeting fragment sizes of 400–600 bp using BluePippin (Sage Science). The size distributions and DNA concentrations of the libraries were evaluated on an Agilent Bioanalyzer High Sensitivity DNA Chip and the Qubit DNA Assay Kit on a Qubit 2.0 Fluorometer (Life Technologies, Carlsbad, CA, United States). DNA concentrations were measured using quantitative PCR prior to cluster generation on the Illumina cBotwhich is a highly advanced automated clonal amplification system at the core of the Illumina sequencing process (Illumina, USA). Finally, single-read sequencing was performed on the Illumina Novaseq 6000 platform (Illumina, USA) for 120 cycles according to Illumina recommendations and included a 1% Illumina PhiX library as an internal control.

### 2.4. Data processing and analysis

#### 2.4.1. GBS data denovo analysis and SNP calling

After demultiplexing the barcoded sequences using the CASAVA pipeline v.1.8 (Illumina, Inc.), the GBS data were processed using GBS2popgen.sh (available on GitHub at: https://github.com/Kferchichi/Automated_stacks), a bash script based on the Stacks pipeline v2.62 (Catchen et et al., 2011) which combines all the steps to process the GBS data, including quality control of the processed reads using FASTQC v0.11.9 (Andrews et al., 2010), and removal of adapters using CUTADAPT v1.16 (Martin et al., 2011). Quality control was then performed using FASTQC after read cleaning to remove any adapter contamination. MultiQC v1.0 (Ewels et al., 2016) was used to compare all reads to each other and to check the final quality version. Denovo assembly of GBS data was performed using Stacks (v 2.63). The Stacks pipeline consists of multiple tools, including ustacks, cstacks, sstacks, gstacks, tsv2bam, and populations, which are utilized in the specific order during data analysis. The ustacks tool was employed with default parameters to create denovo loci for each sample. The cstacks tool was then used to generate a catalogue of loci from the samples provided in the population map, allowing six mismatches between sample loci (*-n 6*) and using a k-mer size of 15 (*--k_len 15*). sstacks was employed to match samples in the population map to the catalogue created in the previous step. tsv2bam was utilized to organize the data by locus rather than by sample, and then gstacks was used to align reads per sample and call variant sites.

For further analysis, the population tool was used to calculate haplotype and SNP-based F-statistics and divergence from Hardy-Weinberg equilibrium for each locus, including only loci, which are present in at least 65 % of the samples within a population (-r 0.65) --fstats –hwe -t 8), to generate SNPs and haplotypes in the variant call format (VCF), which facilitated efficient analysis and identification of informative variants important for downstream population genetic analyses. To refine the dataset, the original VCF file was filtered in VCFtools v4.2 (Danecek et al., 2011) to remove monomorphic SNPs, SNPs with minor allele frequency (MAF) less than 0.05, and a missing data parameter of 0.5 was applied. After quality control, the retained SNPs were processed by the Populations tool to create a new SNPs dataset (genepop=output results in GenePop format; structure=output results in Structure format) needed to determine the population structure.

#### 2.4.2. Within-populations nucleotide diversity and individual heterozygosity

Population genetics software was used to compute estimates of variation among populations. The Populations tool in Stacks provided measures of genetic diversity (π), heterozygosity and of the inbreeding coefficient F_IS_ (Supplementary Table S1). Additionally, we calculated the fixation index F_ST_ for each pair of populations. Based on the datasets before and after filtering, and for comparability between populations, we calculated the normalized numbers of polymorphic sites for each population: we divided the total number of polymorphic sites by the *(2n-1)*-st harmonic number, where *n* is the number of diploid individuals sampled per population. Besides providing comparability, they also represent Watterson’s estimate of the scaled mutation rate *θ_W_=4Nμ* for each population, where *N* is effective population size and *μ* is here the mutation rate per genome per generation. (Supplementary Table S4).

#### 2.4.3. Population structure analysis

To investigate the population structure, a principal component analysis (PCA) was performed using PLINK v1.9 (Chang et al., 2015). The scores of the samples for the first five principal components, along with the eigenvalues of the principal components, were computed and then plotted using ggplot2 v.3.4.2 (Wickham et al., 2016). The model-based clustering software STRUCTURE v2.3.4 (Pritchard et al., 2000) was used to investigate population structure and estimate the allele-frequency divergence among assumed populations. The degree of population substructure was investigated without a priori information other than genotype data. The structure file format output by the Stacks pipeline was adapted according to the requirements of the software. The analysis was performed on 183 individuals of *H. chilensis* with the full dataset (SNP loci in structure format). A series of K = 2,…,6 clusters were tested in ten independent runs. The admixture model with correlated allele frequencies was adopted with each simulation set to a 100,000 burn-in period and 500,000 Markov chain Monte Carlo (MCMC) repetitions. The Q matrices of the estimated membership produced from 10 runs for each cluster were averaged using CLUMPP v.1.1.2b (Mattias et al., 2007) and applying 100,000 repetitions to calculate the mean of these matrices across all replicates. Based on the EVANNO method, STRUCTURE HARVESTER v0.6.94 (Earl et al., 2012) was used to assess and visualize the likelihood values across 10 iterations for each of the six *K* clusters, to more easily identify the optimal number of genetic groups that best describe our data. Finally, Distruct v.1.1 (Rosenberg et al., 2004) was used to perform a customized display of the results produced by CLUMPP in a bar plot for each choice of *K*.

In addition to the default simulation scenario mentioned previously, we conducted a systematic comparison of k-means clustering and the model-based clustering methods in STRUCTURE using the same dataset and parameters. We examined three different scenarios of clustering, including two runs without admixture using different parameters for allele frequencies correlation (correlated and uncorrelated allele frequencies), and one run with admixture and uncorrelated allele frequencies to explore the impact of allele frequencies correlation in the population structure analysis. (Supplementary Figure S2)

#### 2.4.4. Geographical, topographical and precipitation distribution of the different sites

We used the Copernicus GLO-30 digital elevation model (DEM) to illustrate the location of *Huidrobia chilensis* sampling sites and to analyze their geographic and topographic distribution in comparison with the genetic data.

Precipitation for the Atacama Desert from January 2001 to December 2021 was obtained from the Integrated Multisatellite Retrievals for the global precipitation measurements GPM (IMERG; https://gpm.nasa.gov/data/directory). Precipitation in GPM IMERG is estimated using a quasi-Lagrangian time interpolation algorithm. This algorithm intercalibrates, merges, and interpolates multi-satellite microwave precipitation estimates, resulting in a spatial resolution of 10 km. At each sampling station, we calculate the mean precipitation from IMERG using an inverse distance-weighted average over the four nearest grid points. Annual sums of precipitation were calculated per site. Then we determined the MAP per population which was used to create the precipitation difference matrix (Supplementary Table S1).

#### 2.4.5. Genetic distance vs geographical distance

Regression analysis was conducted to examine the correlation between the genetic similarities of population pairs and the corresponding distances separating their geographic locations. This approach allowed us to analyze the spatial genetic structure of the populations and to identify the potential influence of geographic barriers on genetic differentiation. The pairwise genetic distance matrices provided information on the genetic relatedness between individuals, while the GPS matrices resulted in geographic distances between sampling sites. Pairwise genetic distance matrices were generated using the Stacks Populations tool and manually formatted. The Geographic Distance Matrix Generator v.1.2.3 was utilized to calculate the GPS matrix based on longitude and latitude coordinates. This GPS matrix was then used for visualization in R with the package pheatmap v.1.0.12 (Kolde et al., 2019). The resulting heatmaps visualized the pairwise genetic distance and geographic distance matrices, with clustering indicating genetic and geographic similarity between individuals or groups.

Vegan v.2.6.4 (Oksanen et al., 2022), ape v.5.6-2 (Paradis & Schliep, 2019) and sna v.2.7-1 (Butts, 2016) packages in R, were required to determine the Mantel test, calculate the correlation coefficient *r* as well as p-value and plot the correlation between genetic distances and each of the geographic distance, absolute elevation differences and absolute rainfall differences. The Wilcoxon signed-rank test was implemented to detect significant differences between populations in the gene diversity estimates for both geographic and genetic datasets, using the stats v.3.6.2 core package of R (R Core Team, 2023).

## 3. Results

### 3.1. SNPs discovery and data visualization

Of the 186 genotyped individuals, three had to be excluded from further analysis due to poor read quality. The first dataset contained 597,144 loci with an average length of 101 bp per locus. For a locus to be further processed, it had to be present in all populations and in at least 65 % of all individuals in each population, leading to 21,144 loci with a mean length of 111 bp and a total of 55,971 variants after filtering. For PCA and population structure analysis, we also required that the minor allele frequency (MAF) was at least 5 %. This filtering criterion was met by 17,371 SNPs. The first three principal components account for almost 70 % of the variability in the data with the first component explaining 36.2 % of the total variance, followed by the second (20.7 %), third (12.1 %), fourth (10.8 %) and the fifth (4.18 %). Looking at the first two principal components, it is evident that the northern population 807 (Fig. 3A) is separated from the southern populations (Fig. 3). The remaining populations are less clearly separated, forming two less distinct clusters. The plot shows that populations 808 and 809, represented by blue color, are tightly clustered together in the upper right quadrant, along with populations 803, 804, 805, 806, and 810 represented by red color. In contrast, also the red colored populations 800, 801, and 802, are clustered together in the lower right quadrant.

**Fig. 3.**
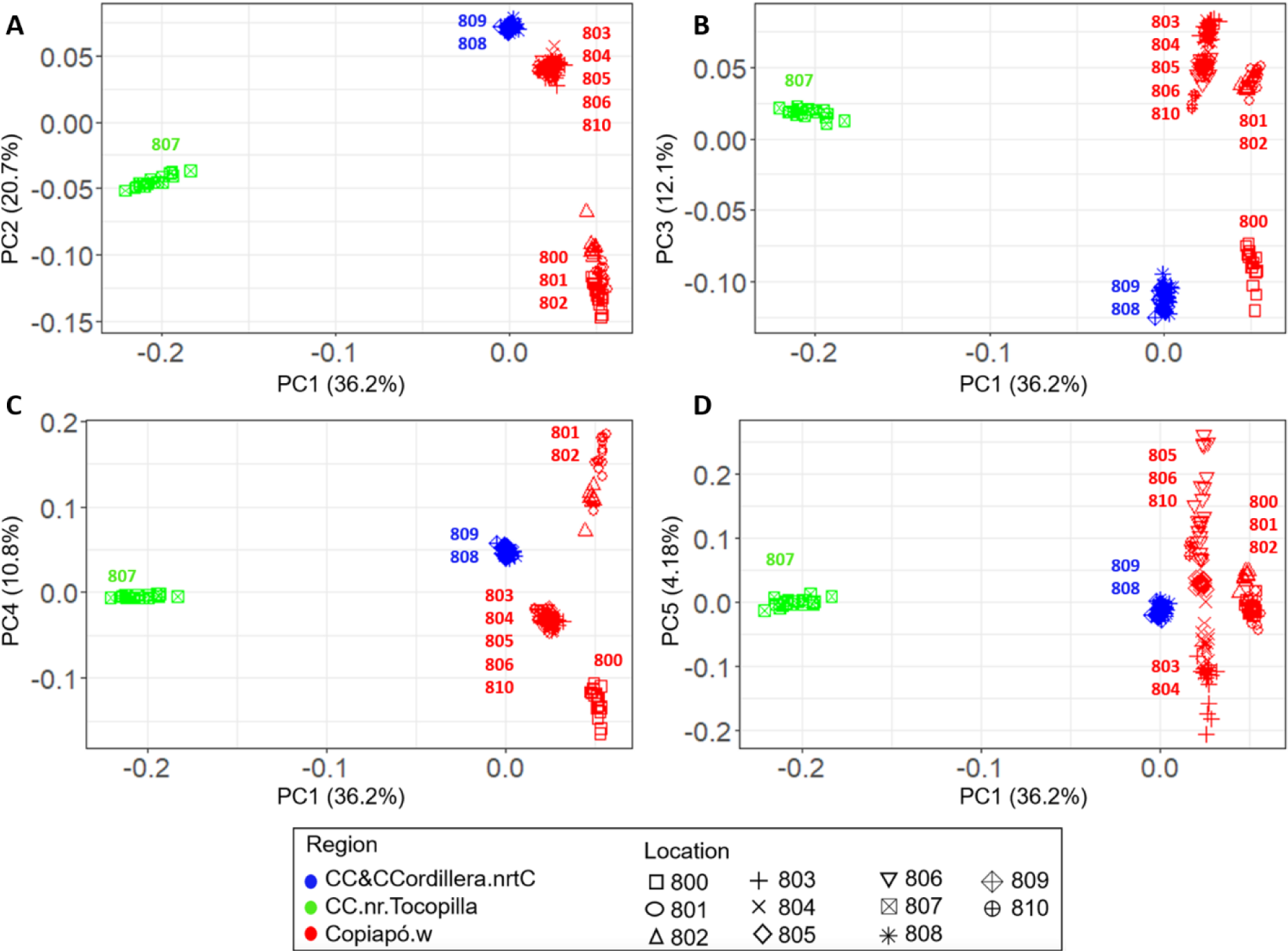
PCA analysis of the broad-scale structure in H. chilensis based on GBS data. PCA was applied to the combined data from all individuals. Green color codes for Coastal Cliff near Tocopilla (**CC.nr.Tocopilla)**; Blue codes for Coastal Cliff and Coastal Cordillera north of Chañaral (**CC&CCordillera.nrtC)**; Red codes for Copiapó Watershed (**Copiapó. w**). **(A)** PC1 and PC2; **(B)** PC1 and PC3; **(C)** PC1 and PC4; **(D)** PC1 and PC5. Clearly visible is separation of population 807 from all others.

### 3.2. Genetic diversity and heterozygosity among populations

Using the normalized number of polymorphic sites, we could show that the northern population 807 has the lowest number of polymorphic sites in both datasets (Fig. 4) (1760 in the filtered dataset and 2827 in the non-filtered dataset), followed by three southern populations 800, 801 and 802 with 2493/3497, 2523/3502 and 2812/3771 respectively and for the filtered/non-filtered datasets.

**Fig. 4.**
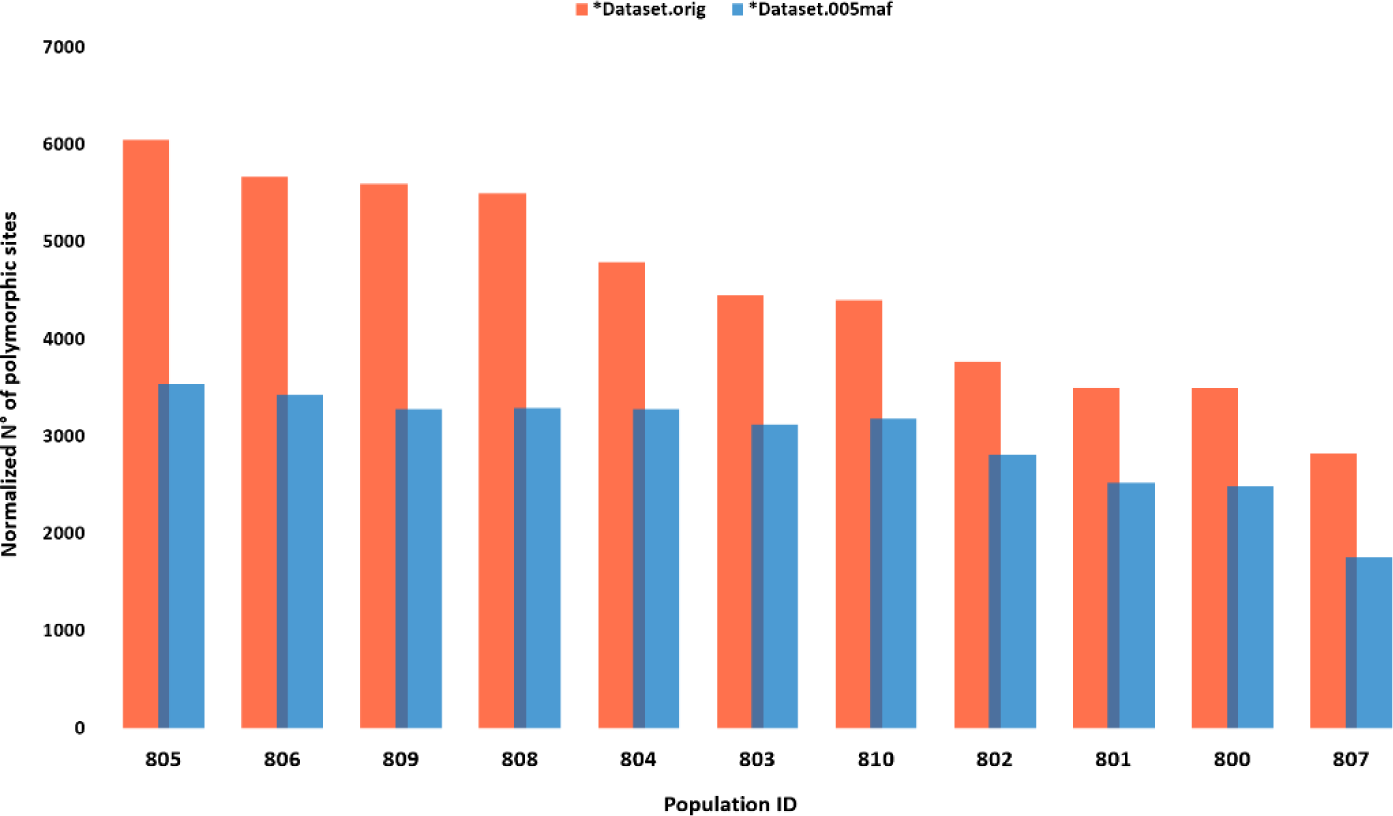
Histogram of the normalized number of polymorphic sites for all populations. Blue color is only those polymorphisms (SNPs) with a pooled (i.e., data from all populations are pooled) minor allele frequency of 5% are considered. Orange color codes for all SNPs are considered. Normalization is done by dividing the raw counts by the harmonic number *H(2n-1)*, where *n* is the number of (diploid) individuals sampled per population.

Based on the filtered dataset, we found that Population 807, consisting of 20 individuals, shows the highest number of private alleles (452) compared to the other populations with the same sample size (Supplementary Table S5). Populations 800 had only 50 private alleles. Only one private allele each were found in populations 809 and 803 (19 individuals). No private alleles were identified in populations 803 (19 individuals), 804 (17 individuals), 805 (20 individuals), 806 (20 individuals) and 810 (4 individuals). The number of polymorphic sites varied from 11,413 in population 810 (with only 4 individuals) to 25,718 in population 805, with 20 individuals (Table 2). However, population 810 has the lowest percentage of polymorphic sites (0.673 %), followed by populations 807 (0.676 %) and 802 (0.689%) (Table 2).

**Table. 1.**
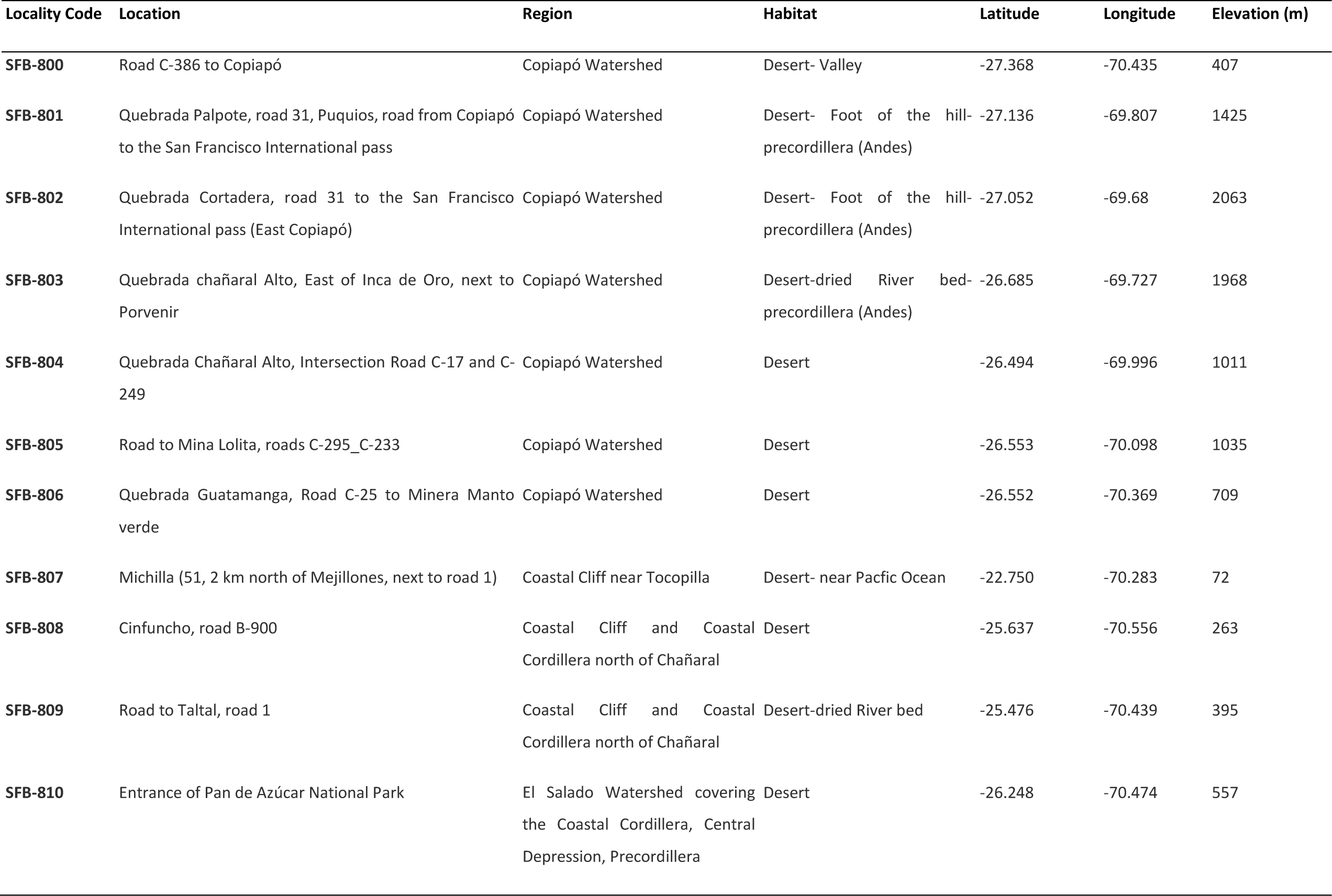
Sampling data of *Huidobria chilensis* collected from 11 different locations; SFB indicates the project’s ID and the numbers from 800 to 810 refers to the different stations.

**Table. 2.**
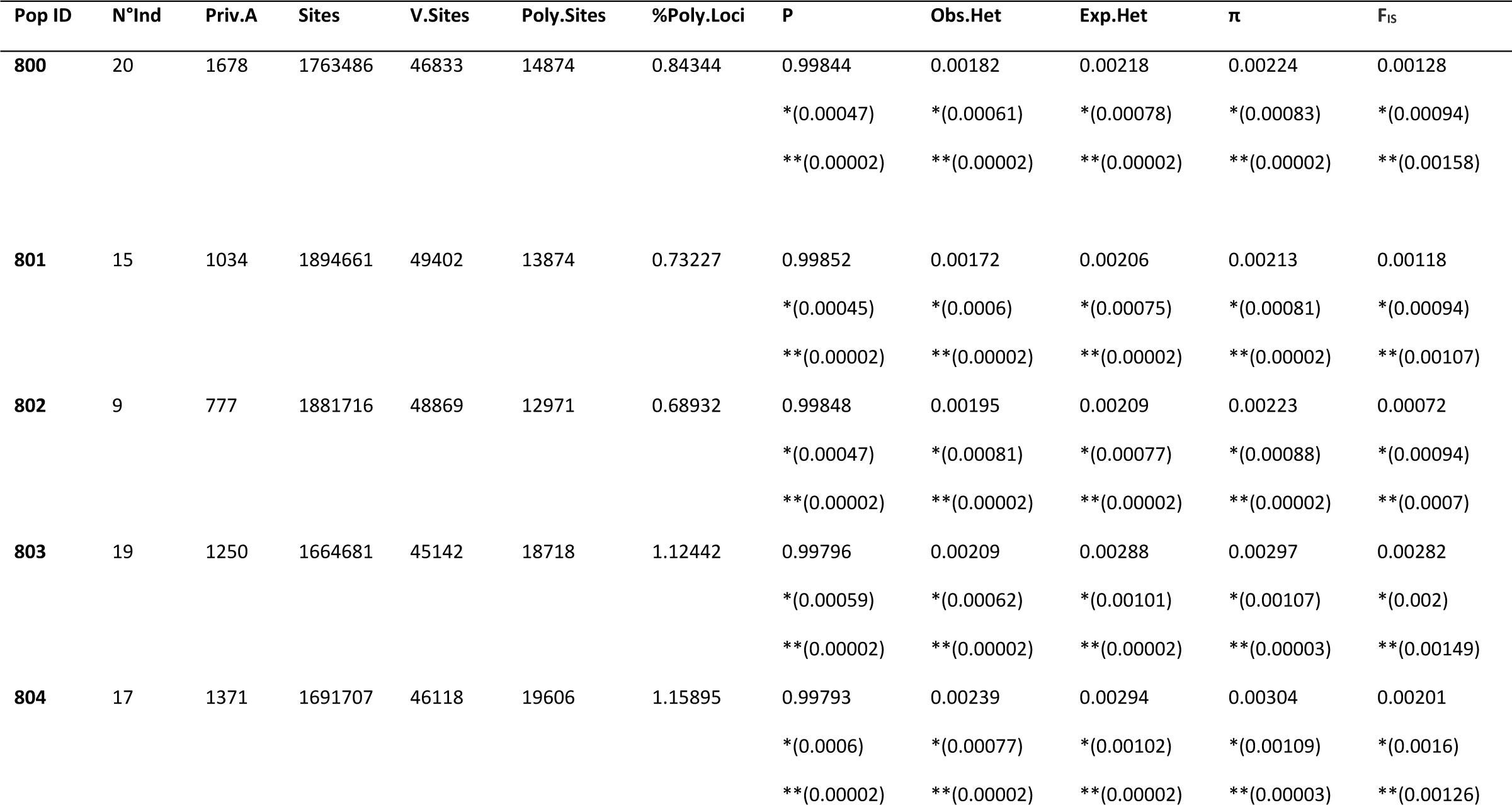
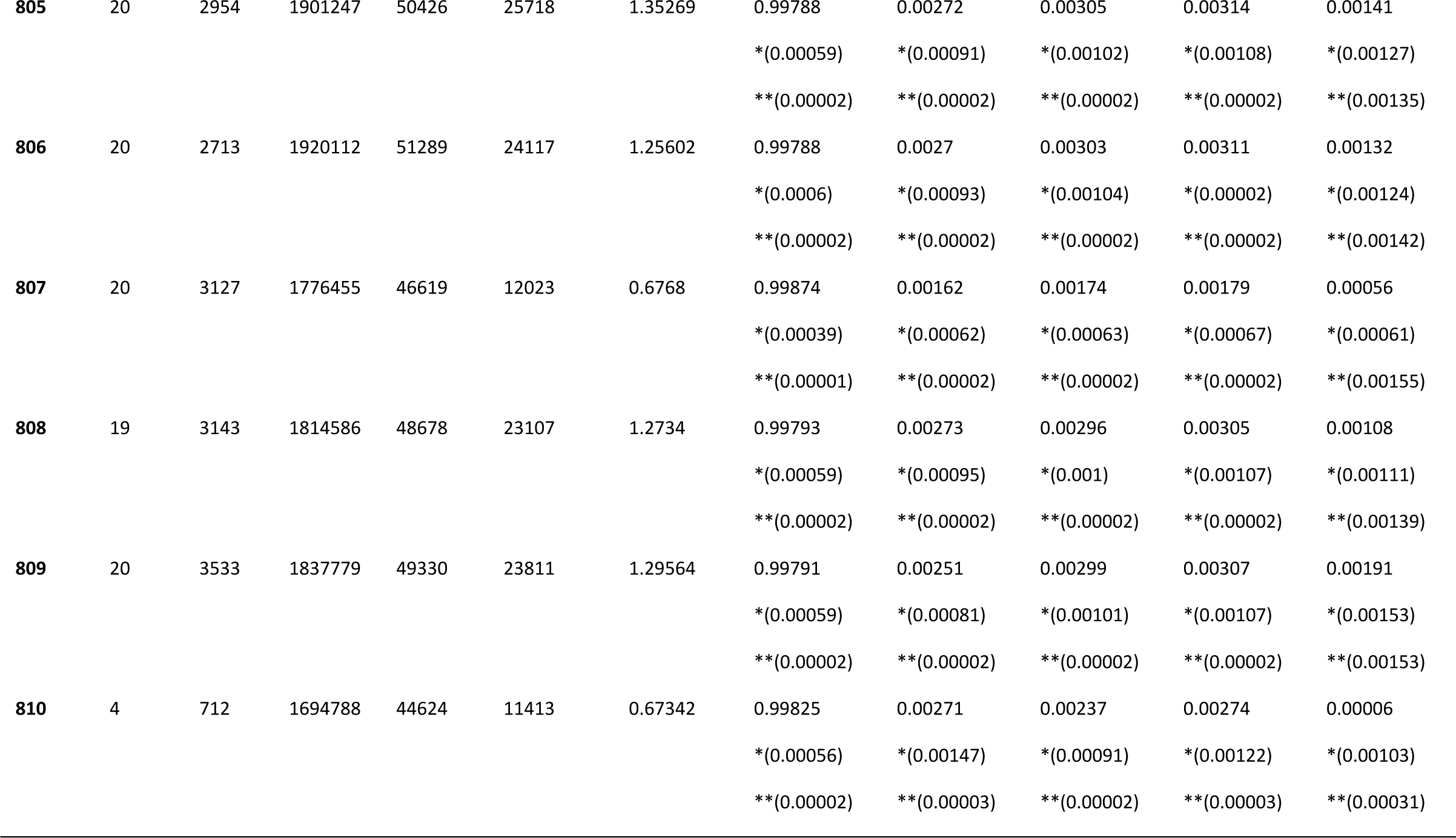
Table summarizing statistics provided by “Populations” tool of stacks. Pop ID: Populations identifiers; N° Ind: the studied number of individual for each population; Priv.A: number of private alleles in this population; Sites: Number of sites in the dataset; V.Sites: Number of variant sites; Poly.Sites: Number of polymorphic sites per population; %Poly.Loci: Percentage of polymorphic loci; P: Mean frequency of the most frequent alles in each locus for this population; Obs.Het: Observed heterozygosity; Exp.Het: Expected heterozygosity; π: nucleotide diversity;; F_IS_: inbreeding coefficient; *r: Variance; **: Standard error.

Accounting for the different sample’s sizes and the different numbers of genotyped loci among the different populations, the estimated nucleotide diversity π was relatively similar in all populations: it ranges from 0.00213 in population 801 to 0.00314 in population 805, except for population 807 in the north, which has a π of 0.00179 (Table 2). Both observed and (at Hardy-Weinberg equilibrium) expected hterozygosity are lowest and highest in populations 807 and 805, respectively. Wright’s F_IS_, which measures the deviation from Hardy-Weinberg equilibrium within a population, is close to zero and ranges from 0.00006 in population 810 to 0.00282 in population 803 (Table 2). Generally, positive F_IS_ may be caused by inbreeding or selfing, and negative F_IS_ indicates an excess of heterozygotes. When seen on a genome-wide scale, this could be a signature of some degree of clonal reproduction. (Table 2).

### 3.3. Population structure

The population structure of 183 *H. chilensis* individuals was assessed on the basis of 17,344 loci using the clustering program STRUCTURE with an admixture model. Identification of the optimal number of sub-populations (*K*) was based on maximum likelihood and the so-called “delta *K*” values, provided as output from STRUCTURE. Our dataset was found to be best explained by assuming three main genetic groups (*K* = 3). All other K values examined (*K* up to 6) resulted in lower delta *K* values (Supplementary Table S8).

When choosing *K* = 2, the population assignment separates the northern population (807) from all others (Fig. 5C). We could also see that the southern and northern gene pools were clearly distinguished, with a low admixture observed in 810, 809 and 808 with respect to 807; which is persistent at *K* = 3.

**Fig. 5.**
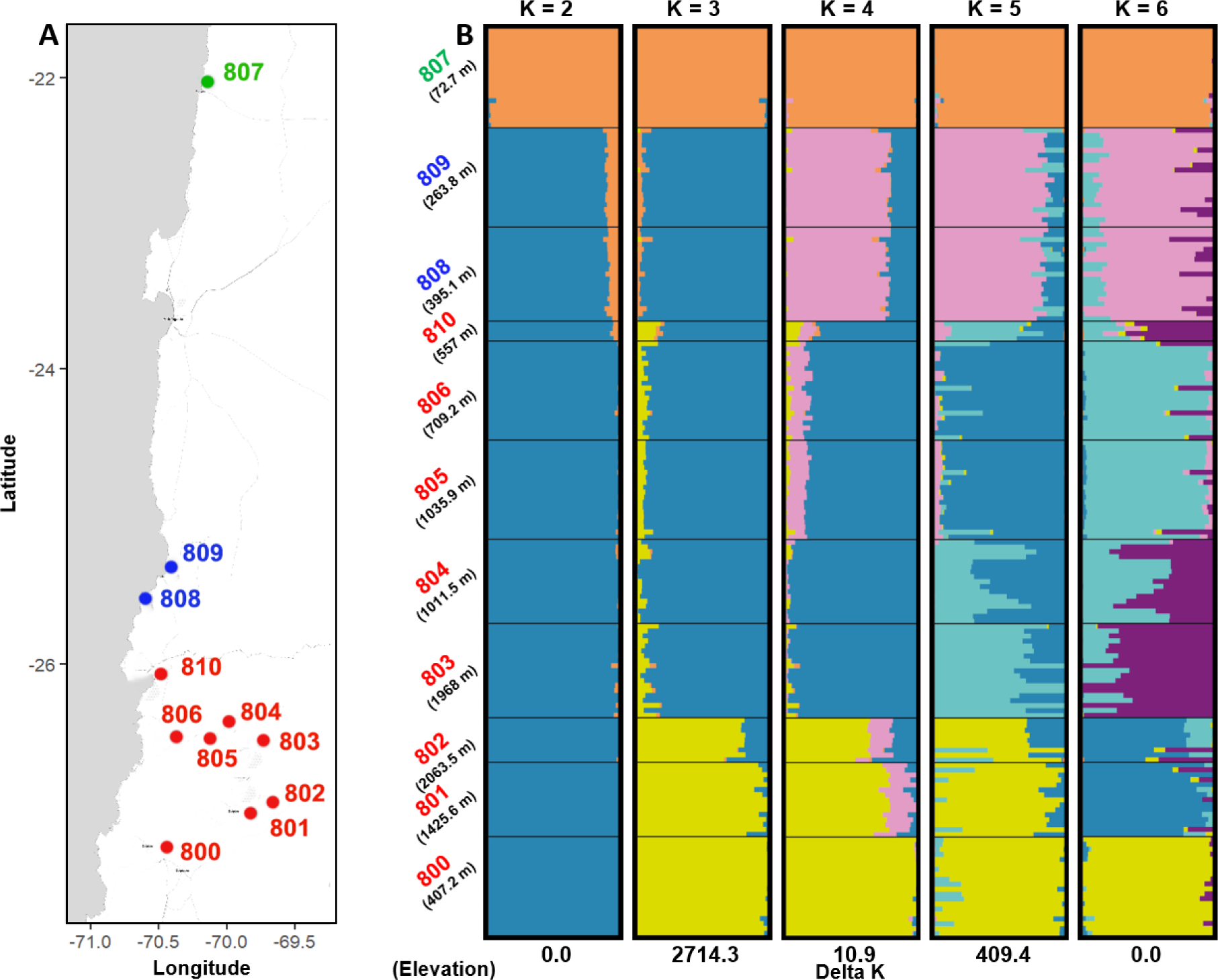
Population STRUCTURE of eleven populations of *Huidobria chilensis* using admixture based-model Structure software with correlated allele frequencies. Results were averaged using CLUMPP software and plotted by Distruct software. **(A)** Map generated using ggmap showing the sampling sites; **(B)** Clustering of 183 *H. chilensis* individuals presented by different colors i.e., 1. Orange, 2. Blue, 3. Yellow, 4. Pink, 5. Light blue and 6. Dark purple inferred by Structure analysis for estimating the mean likelihoods of each number of genetic clusters, the given values with population number on the left side of the plots are the elevation data corresponding to each site. Each bar representing an individual, where different colors and the length of each segment representing the estimated membership coefficient to each of the genetic clusters using the Q matrices generated from the STRUCTURE simulations. K suggested a division into two or six genetic clusters. 0, 1.3, 0.5, 17.4 and 0 are respectively delta Ks for the clustering K2 to K6. Three different regions are coded as follow: Green (for site 807): Coastal Cliff near Tocopilla; Blue (for sites 808-809): Coastal Cliff and Coastal Cordillera north of Chañaral; **Red** (for sites 800-806 and 810): Copiapó Watershed.

For *K* = 3, the subpopulation excluding 807 was further divided into three recombined regions: a small part from Copiapó including only 800 to 802 (catchment 1), the rest part of Copiapó including populations 803 to population 806 with 810 (catchment 2), and the Coastal Cordillera north of Chañaral including 808 and 809 respectively (Fig. 5B). The samples from populations 800 to 802 show admixture, partly assigned to a third ancestral population, partly sharing ancestry with populations 803 to 806 with 810, and a very small proportion with both populations 808 to 809. In addition, nearly 89 % of all individuals (163 out of 183) showed no shared ancestry and were assigned to exactly one population each. The remaining 11 % of the individuals were considered to be of mixed ancestry. In particular, populations 800, 801, and 802 exhibited strong evidence of heterogeneity. As K increased, more and more populations showed mixed ancestry.

With *K* = 4, there are four clusters corresponding to three regions in Copiapó, including the population 810, while 808 and 809 were clearly separated from their previous cluster to create a cluster of their own, 807 was still assigned to one separate cluster. The populations 800, 801 and 802 form a joint ancestry group, with more admixture observed in 801, 801 and 802 with the other populations except 807. However, 808 and 809 are shown as admixed. Part of the alleles are assigned to a separate cluster; another part is shared with populations 801 to 806 (Fig. 5B). The northern cluster consisted of 44 individuals that belong to three different populations 800, 801 and 802 located in the same geographical region. Another cluster (blue color, Fig. 5B) consisted of 80 individuals from 5 different populations 803, 804, 805, 806 and 810, located in two different regions. Further cluster was composed of 39 individuals belonging to two populations located in the same region and very close to each other. The fourth cluster (orange color) contained individuals from one population (807) in a geographically separate area.

The STRUCTURE bar graph (Fig. 5C) offers valuable insight into the degree of admixture present in the studied material. For *K* = 5, approximately 170 individuals were found to be of mixed ancestry. Populations 800, 801, and 802 showed more admixture, than 805 and 806. Lowest admixture was observed in populations 808 and 809. For *K* = 6, population 810, located between the northern and central regions, is assigned to a distinct cluster.

The fixation index F_ST_ measures population differentiation. We find higher differentiation between populations 800, 801 and 802 (average 0.10) than between populations 803, 804, 805 and 806 (average 0.033). Both clusters belong to the southern cluster (Copiapó). Also mean F_ST_ between 810 and 800–802 (0.146) is higher than the mean F_ST_ between 810 and 803–806 (0.052). Mean F_ST_ is also very low (0.03) between populations 808 and 809 from the central geographic cluster (Coastal Cordillera north of Chañaral), while mean F_ST_ between these two populations and the northern population 807 (near Tocopilla) was quite high (0.18), similar to the mean between population 807 and 805–806 from the south, and slightly lower than the mean between population 807 and 803– 804 (0.19) from the same southern geographic cluster. The highest F_ST_ values were identified in all pairwise comparisons containing 807. The highest F_ST_ (0.26) is identified in the pair 807–801, followed by 807–802 (0.26), 807–800 (0.25) and 807–810 (0.25). (Fig. 6A).

**Fig. 6.**
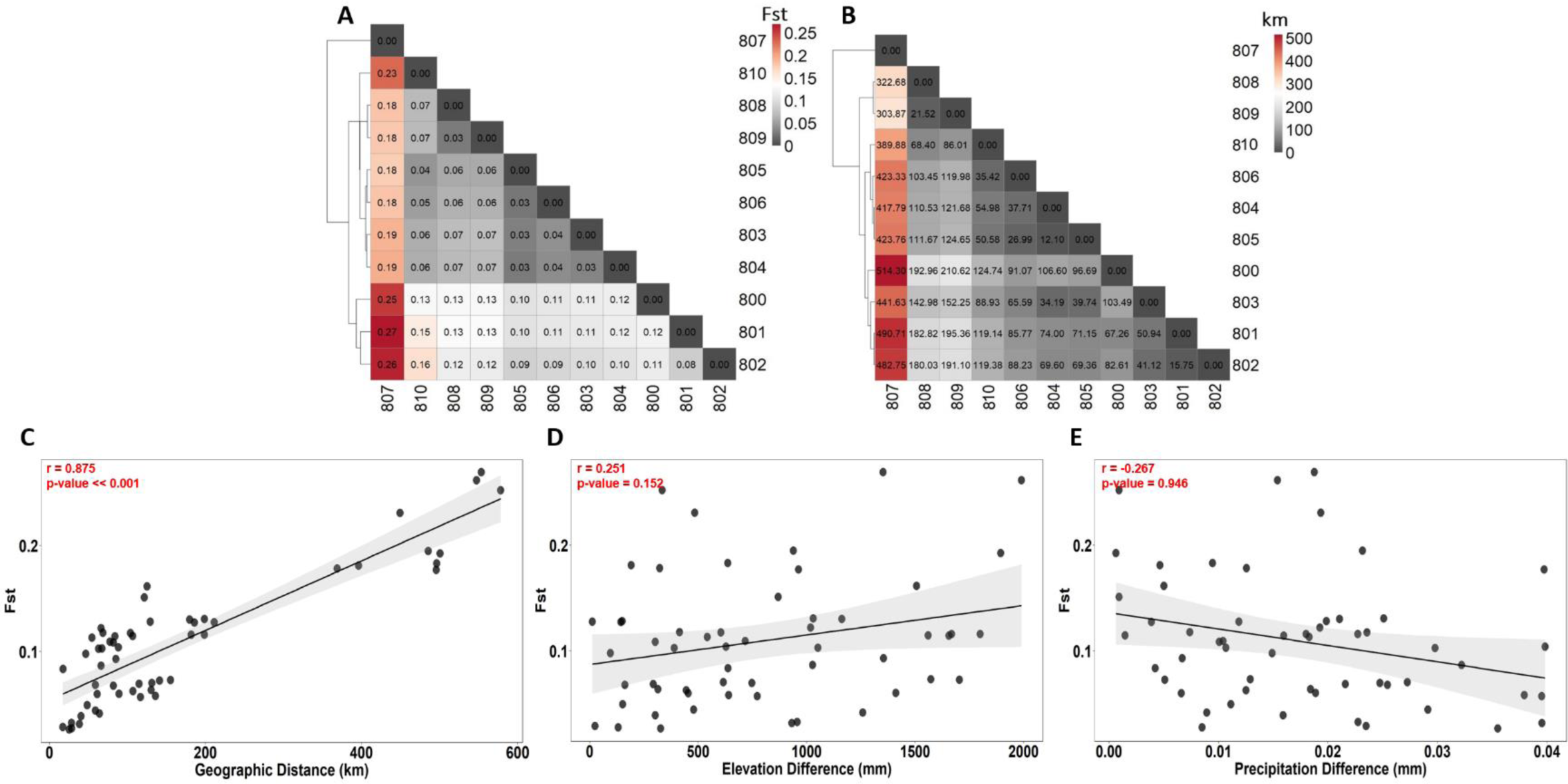
Correlation between genetic, geographic and environmental variables in 11 *Huidobria chilensis*’s populations. **(A)** Heatmap of pairwise F_ST_ used as a genetic assignment baseline; Dendrogram clusters the populations according their F_ST_ correlation. **(B)** Heatmap of 10 pairwise distances in km based on GPS coordinates; Dendrogram clusters the populations according their geographic distance’s correlation. Dark gray cells denote lower pairwise F_ST_ estimates and greater genetic similarity among populations, salmon, orange and red cells denote higher F_ST_ and greater genetic divergence; (**C)** Scatterplot showing a linear correlation between F_ST_ and geographic distances of 11 populations of *Huidobria chilensis*; r denotes Pearson’s correlation coefficient with a significant p-value (Mantel’s and rank correlation test were done); (**D)** Scatterplot showing a linear correlation between F_ST_ and elevation difference. No significant correlation is observed; (**E)** Scatterplot showing a linear correlation between F_ST_ and precipitation. Precipitation differences are calculated from yearly precipitation values and averaged over a period of 20 years (2001-2021). A negative correlation is observed (Mantel statistic *r* = -0.267, Significance p-value = 0.946).

### 3.4. Abiotic factors and genetic diversity

According to the Mantel and ranking tests (p-value << 0.01), genetic differentiation correlates positively with geographic distance (*r* = 0.875) (Fig. 6C). Population 807 lies at a distance of 514.30 km, 490.71 km, 482.75 km and 389.88 km from populations 800, 801, 802 and 810, respectively. In addition, based on the IMERG data from weather stations located near the current sampling sites and collected over the past 20 years showing the difference in long-term average precipitation between any two stations, the Mantel test performed on the rainfall and the F_ST_ matrices yielded a negative correlation explained by *r* = -0.267 and P > 0.01 (Fig. 6E).

Similarly, the Mantel test did not establish a significant correlation between F_ST_ and elevation differences in every pairwise populations. Interestingly, the trend shows even a low *r* = 251 with P > 0.01 (Fig. 6D).

## 4. Discussion

We investigated three distinct population clusters of *H. chilensis* from three sampling regions of the Atacama Desert. Patterns of genetic differentiation were analyzed with the program STRUCTURE v2.3.4 (Falush et al., 2003), and with principal component analysis (PCA) (Figs. 3, 5) based on the Evanno method (Supplementary Figure S1). The results revealed that *K* = 3 is the number of genetic clusters which best explains the geographic distribution of *H. chilensis.* The results show a relatively low genetic diversity within populations. Both the observed heterozygosity (H.obs) and genetic diversity π range between 0.002 and 0.003, and the observed F_ST_ values (F_ST_ ≤ 0.268) are slightly higher than those found among 21 populations of the Chilean endemic species *Tillandsia landbeckii* (F_ST_ ≤ 0.222; Merklinger et al., 2020). For the sister species *H. fruticosa* Merklinger et al. (in prep.) reported F_ST_ values of up to 0.35, indicating a somewhat stronger genetic differentiation in that species at a comparable geographic distance.

### 4.1. Genetic differentiation and population structure

The analyses of the population genomic GBS data showed a clear separation of the northern and southern populations with a moderate level of divergence and identified three genetic clusters corresponding to distinct phylogeographic regions. One is located in the coastal cliff region between Antofagasta and Tocopilla (northern population: 807, see Fig. 2, one between the coastal cliff and the Coastal Cordillera further south (“central” populations: 808–809, see Fig. 2), north of Chañaral and the Copiapó watershed area, and one in the southern part of the Copiapó region (southern populations: 800–806, 810, see Fig. 2). The southern populations could be further divided into two valley groups (vg1: 800–802 and vg2: 803–806).

In general, genetic diversity found in the *H. chilensis* populations is moderately low. In the southern populations it is on average 0.00258 (across 8 populations) with little variation among the individual populations. Across the Copiapó watershed cluster of populations 800, 801, 802 it is somewhat lower (0.00211) than across the Chañaral cluster of populations 803, 804, 805, 806 (0.00285). When restricted to the four populations with a uniform sample size of 20 the average is 0.00279, confirming that our π estimates are not suffering from sample size effects. The two coastal populations in the center are also rather uniform in genetic diversity, with an average of 0.00298. The only exception is the isolated northern population (807) with a diversity of only 0.00174. Its low value is likely due to its demography (isolated single population with reduced effective population size and suppressed or limited gene flow from other populations) rather than to a difference in the mutation rate or a stronger selection pressure. Its isolation is also confirmed by its high proportion of private alleles (Supplementary Table S3). Thus, disregarding population 807 for its special situation, genetic diversity in *H. chilensis* at species level seems to be around 0.2% to 0.3%. In all populations we find observed and expected heterozygosities to be very close to each other, without major deviations from Hardy-Weinberg equilibrium. In agreement with this, F_IS_ values (Table 2) are very close to zero and consistently slightly positive. Judging from these there are no evident signatures neither of selfing nor of clonality.

Considering genetic differentiation among populations we calculated F_ST_ for all pairwise comparisons. Except for the comparisons with the northernmost population 807, the F_ST_ values between populations in the central (808–809) and the southern (800–806, 810) regions are low to moderately high (ranging from 0.03 between the neighboring populations 808 and 809, to 0.16 between the distant populations 802 and 810), indicating ongoing gene flow between populations. In contrast, the F_ST_ analysis again confirms the relative isolation of population 807. F_ST_ values range from 0.18 (differentiation between 807 and the two central populations) to 0.26 (between 807 and 802). In addition, the pairwise F_ST_ values between the 11 populations (Supplementary Table S7) in general confirm that genetic substructure in *H. chilensis* is mainly driven by geographic distance.

### 4.2. Spatial patterns of genetic diversity

The topography of the Atacama Desert and the distance of more than 500 km between populations 807 and 800 are limiting factors for the dispersal of this plant. In addition, there seems to be a lack of “stepping stones” (Kimura and Weiss, 1964), which most likely contributes to the isolation of 807. To the best of our knowledge, we could not identify any accessions of *H. chilensis* between the locations of 807 and of 809, resulting in a sampling gap. The genetic data are consistent with such a habitat gap, yet according to data from the Global Biodiversity Information Facility (GBIF.org; accessed 21.04.2023; https://doi.org/10.15468/dl.tfhnmx), few occurrence records in the wider Antofagasta region are documented, leaving an effective gap of about 150 km. Thus, we cannot completely exclude the occurrence of *H. chilensis* in this gap, despite the clear genetic data.

Based on both Mantel and ranking tests (Fig. 6C), we observed a strong correlation (*r* = 0.875) between genetic differentiation and geographic distances, suggesting a pattern of isolation by distance (Fig. 6C). This correlation is higher compared to those reported in *H. fruticosa* (*r* = 0.749; Merklinger et al., in prep) or *Tillandsia landbeckii* (*r* = 0.72; Merklinger et al., 2020). Additionally, we identified some cases of admixture between populations that were not geographically far apart, probably as a result of water flow patterns in the surrounding area. In addition, the Copiapó basin may have admixed populations 800–802 (vg1), but the frequent genetic exchange among individuals in these populations may have occurred because they occupy the same ecological niche and are in close proximity, resulting in cross-pollination. Precipitation patterns and variability, controlled by elevation and glacial-interglacial cycles (McGee, 2020), may have further contributed to the development of distinct groups. In the lowlands, the complex topography and lack of groundwater systems connected to higher elevations that receive sufficient precipitation (Fig. 7) may have restricted the movement and mixing of species.

**Fig. 7.**
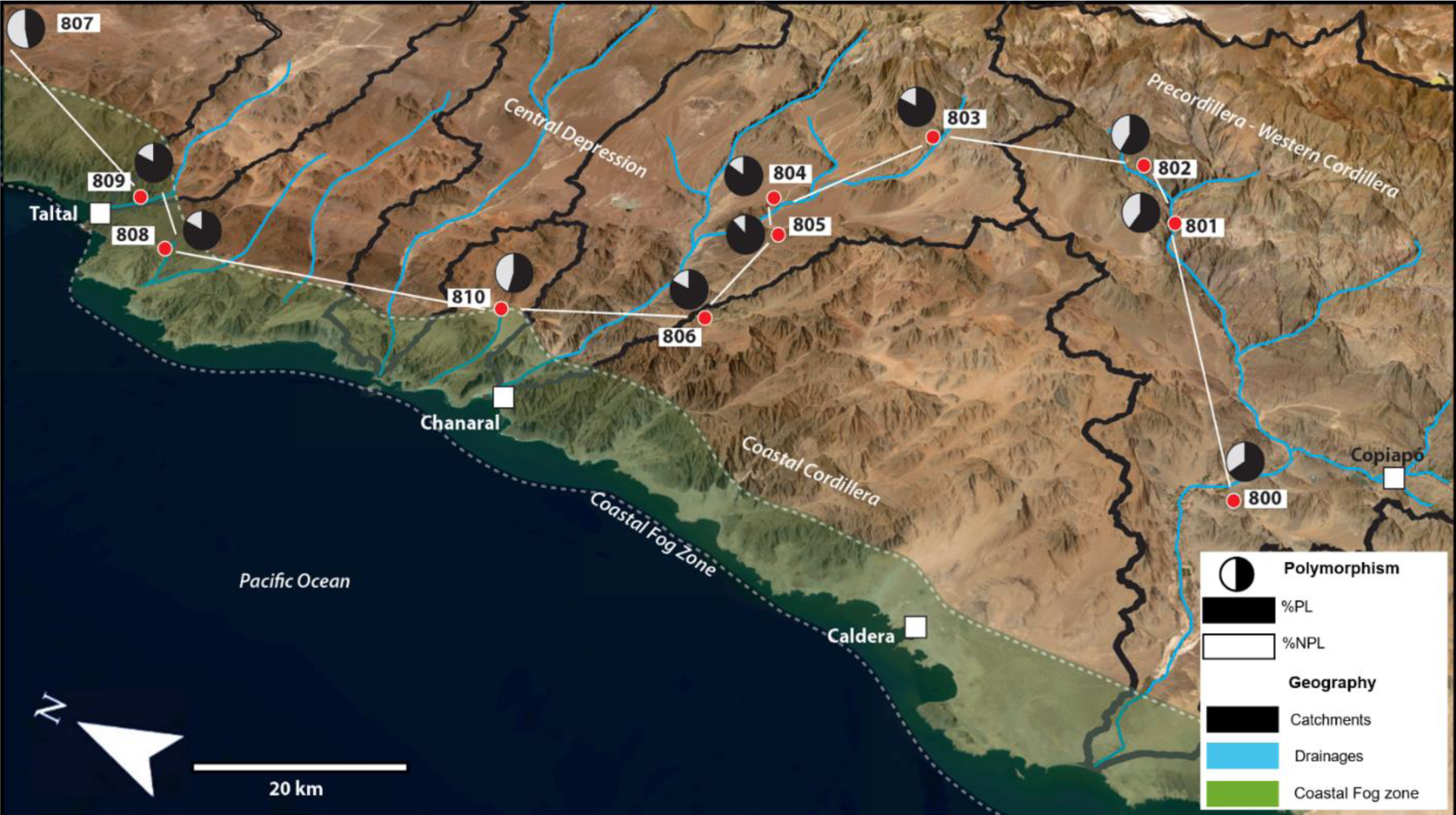
3D Map showing the major environmental and genetic features associated with the sampling sites of *Huidobria chilensis* mapped using ArcGIS Pro v.3.1.0 and Adobe illustrator 2023 software. Drainages (blue), Catchments of the sampled populations (black). Pie charts represents the genetic diversity pattern for each site. Percentages of polymorphic Loci were calculated from the normalized numbers of all sites and polymorphic sites (from the filtered dataset) (Supplementary Table. 2) in every population. The coastal fog zone is only approximately sketched.

We suggest that *H. chilensis*, like *H. fruticosa*, is associated and restricted to larger catchments with sufficient groundwater from the headwaters of the respective catchments, or they thrive exclusively in valleys where sufficient moisture can be collected from local precipitation. Therefore, connectivity and the gene exchange are only possible if the surrounding environmental conditions allow migration between populations. It seems that both topographic and climatic factors may have contributed to the cutoff of migration pathways and the cessation of gene flow and exchange between the two southern valley groups. In other words, the physical characteristics of the highlands, especially and change of temperature by climate deterioration since a few million years, cause the narrowing of potential high altitudes habitats, and consequently a cut-off of migration occurred via the highlands. So, populations are forced to lower altitude habitats suitable to their requirements and environmental limitations, this may have hindered migration and differentiation (Weir, 2006), tending to mask any potential signals of elevation on genetic variation (Sousa and Hey, 2013). Moreover, additional factors, such as soil composition, salinity, frequency and periodicity of precipitation events, drought periods, and fog occurrence and quantity, might have an additional influence and impact in the distribution of organism in these extreme environments.

### 4.3. Demography of Huidobria chilensis in response to Quaternary climatic fluctuations

The distribution patterns and phylogeographic structure of species, especially those that are drought-tolerant and living at high elevations, such as in parts of the Atacama Desert, have been significantly influenced by paleoclimate variability (Ossa et al., 2013). Although some insights have been gained from previous research efforts using molecular clock dating to correlate climate history and landscape evolution with diversification and biogeography, for example in the genus *Cristaria* (Böhnert et al., 2019), a comprehensive understanding of the species’ demographic history has been hindered by limited genetic data and population structure studies. In Merklinger et al. (in prep), they used a new „triple comparison method” to estimate the time of sequence divergence among *H. fruticosa* population clusters. According to their results, the separation of the clusters has taken place about approximately 2 million years ago. Similar age constraints are estimated based on molecular clock data for various genera in the Atacama Desert e.g., *Nolana* (Dillon et al., 2009), *Eulychnia* (Merklinger et al. 2019), *Cristaria* (Böhnert et al. 2022), darkling beetles (Zúñiga-Reinoso et al. in prep., Ragionieri et al., 2023). As we do not have molecular clock information to provide additional insight into the timing and subsequent paleoclimate variability in the past, we assume, based on recent publications from the study area (Böhnert et al., 2020, 2022; Zúñiga-Reinoso and Predel 2019, Zúñiga-Reinoso et al., 2021), that the cutoff of genetic exchange between the two main populations was driven by climate degradation with subsequent changes in environmental boundary conditions, such as temperature decrease, permafrost boundary shift, precipitation reduction and modification, since the onset of the Quaternary and presumably enhanced or intensified since the Mid-Pleistocene transition (Berends et al., 2021), with high-amplitude climate oscillations. Thus, considering the lack of chronological information for *H. chilensis,* however, based on the work performed on its sister species *H. fruticosa*, we aim to fill the gap by relating the Quaternary demographic history of *H. chilensis* to current climatic, environmental, and geographic conditions. This change in boundary conditions, combined with the complex topography, has likely caused the closure or narrowing of potential migration corridors for genetic exchange between populations of *H. chilensis*.

The studied area has a complex topography, although that both high altitude populations (802, 803) are geographically proximal to each other (41.2 km linear distance), a mountain ridge separates both populations. Moreover, we assume that due to the proximity of the population, the precipitation pattern and system is indistinguishable. Therefore, separation of previously connected populations was presumably primarily caused by changing environmental boundary conditions during presumably the Quaternary (Merklinger et al., in prep.), especially temperature reduction and subsequently shifting potential habitats to lower elevations. The latter most likely caused the cutoff of gene flow exchange between these populations. The large topographic fragmentation in the Coastal Cordillera between the two lineages or the main populations (808, 809) can be seen as another cause of isolation and reduced genetic exchange. The rugged terrain between the southern catchment (vg1) and the proximal northern site (807) lacks major drainage systems that are connected to the main source of precipitation originating in the high Andes (Fig. 8), reducing the occurrence of potential stepping stones for infrequent or intermittent gene flow exchange during more suitable environmental conditions.

**Fig. 8.**
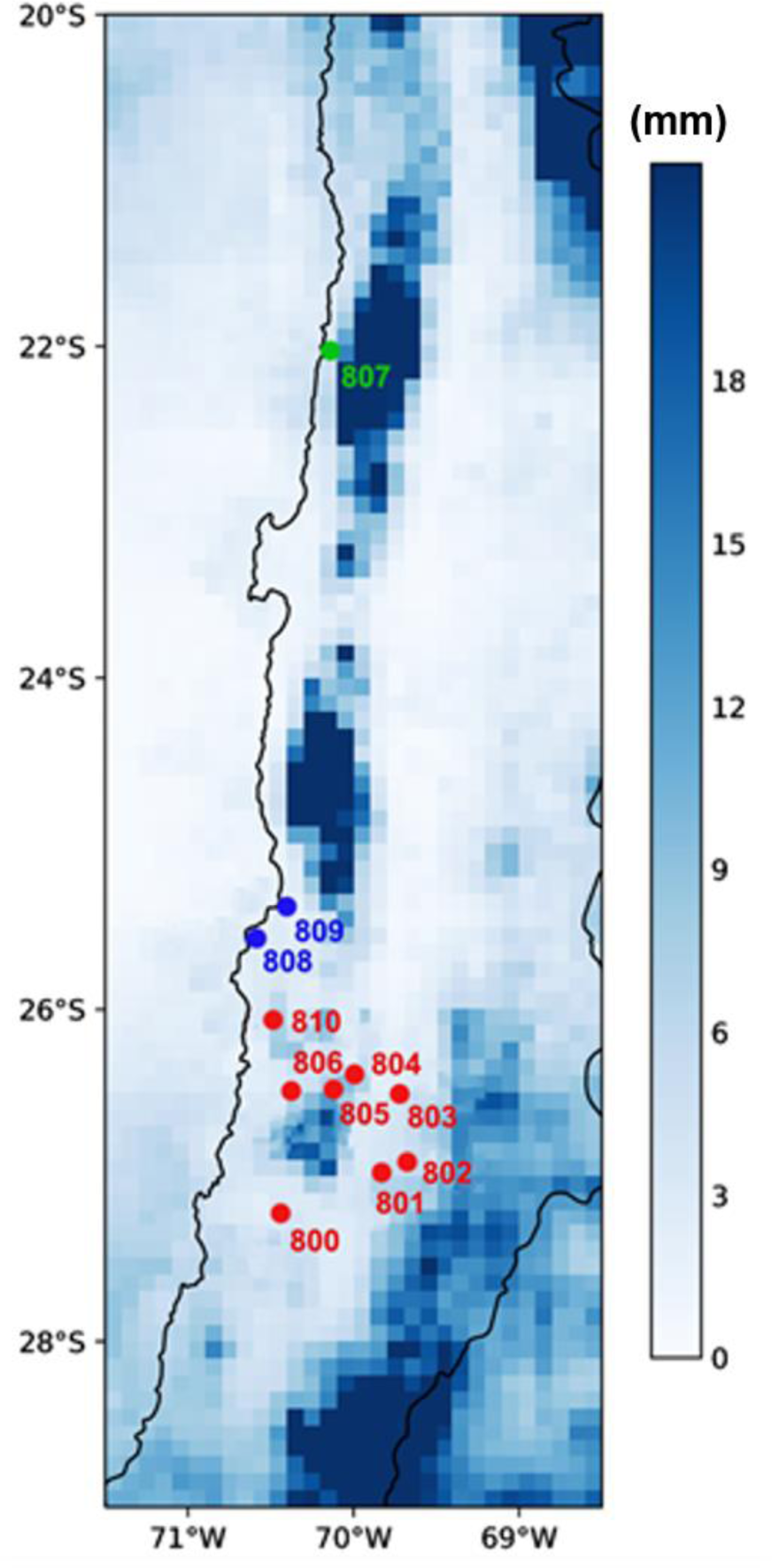
Annual average precipitation in mm per year for the Atacama Desert for January 2001 to December 2021. Data were obtained from the Integrated Multisatellite Retrievals for the **global precipitation measurements** GPM (IMERG; https://gpm.nasa.gov/data/directory). Different regions for the sampling sites are marked in color with green for the Coastal Cliff near Tocopilla, blue for the Coastal Cliff and Coastal Cordillera north of Chañaral, and red for the Copiapó Watershed.

It can be assumed that the recently dispersed populations of *H. chilensis* have been based on either (1) drainages that receive precipitation in adjacent higher terrain and whose groundwater can be accessed by deep roots of *H. chilensis*, or (2) the coastal fog oasis. The area between these two potential sources of precipitation is located in the absolute desert with almost no local precipitation, as such we suggest that drainages connecting and bridging both sources of moisture are the key for migration and genetic exchange. However, the populations 801–805, located at higher elevations, have an average distance of 47.8 km between them, which is shorter than between populations 807, 808, 809 and 800, located at the lowest elevations, which have an average distance of about 261 km, 141.7 km (808, 809 and 800), respectively. Therefore, based on the elevation and rainfall fluctuations data (Supplementary Table S1 & S2), these potential forcing factors were not identified as the main driver for large-scale spatial genetic structure in this species (Fig. 6D, E). The latter could be due to the limited rainfall fluctuation within the study area or other confounding factors, such as the remote location of climate stations from the collection sites (Schween et al., 2020), which could affect the detection of precipitation data. We suggest that the local precipitation in the lowlands may not be sufficient to drive migration and expansion of *H. chilensis* (Fig. 6E). Expansion and connectivity between the southern and central population of *H. chilensis* might be further impeded by the rocky and sandy habitat which could prevent migration and hybridization via a coastal corridor caused by coastal fog along fog oases. Populations 807, 808, and 809 are likely to be exposed to fog, and/or sustain due to fog availability, indicates that genetic diversity in this studied area could be influenced by fog events, which is also supported by Mörchen et al. (2021), based the on n-alkane distribution in fog-fed plants at the margins of the Atacama Desert.

Except for the two high elevation and proximal populations (802, 803) with clear indication of an altitudinal influence on the geneflow cutoff, the elevation range within the study area may not be large enough to cause any forcing on geneflow exchange and diversity. Additionally, it may be explained by the fact that the large elevation differences between specific populations are found among relatively closely located mountain populations (800-805), while the coastal populations are at similar elevation (800 and 807-809) however great distances.

Additionally, the distribution of accessions is likely influenced by precipitation in the higher elevation catchments, which receive more rainfall and may create a topographically top-down controlled system (Ragettli et al., 2016; Vuille, 2014), allowing the survival of *H. chilensis* in the more arid to hyper-arid part of the Atacama Desert ‘lowlands. A down-top controlled system might be at play This explaining the more genetically proximal distance of the northern populations, driven by the occurrence of coastal fog oasis and potential migration pathways along the coast to the north or south. However, it is interesting to note that *H. fruticosa*, which occurs in similar habitats in the Coastal Cordillera as well as on the western slopes of the Andes north of the *H. chilensis* distribution, is reported to occur continuously up to the northern vicinity of the central populations of *H. chilensis* (Merklinger et al., in prep.). This suggests a distributional boundary for *H. chilensis*, due to a yet unknown change in environmental factors, that is not currently understood. This boundary presumably corresponds to a change in annual rainfall (Supplementary Figure S3), but more detailed analyses are needed.

## 5. Conclusion

In this study, we focused on how environmental diversity in the Atacama Desert landscape affects gene flow and genetic differentiation in *H. chilensis* as a result of geographic isolation. Analysis of the GBS data revealed three main findings. First, the southern geographic cluster showed substantial genetic diversity, suggesting that the sexually reproducing individuals in this region likely belong to the oldest populations in the Atacama Desert of Chile. Secondly, greater distances between locations led to increased genetic differentiation, suggesting that migration and gene flow occurred in the past, which are now disturbed or interrupted. Current geographical barriers contribute to the preservation of these genetic patterns, particularly between the northern and southern clusters, while the northern population may be a relict of the past. Finally, region-specific factors, rather than physical barriers and precipitation, were found to drive genetic divergence, particularly between the population 807 and all others. Hence, by studying further environmental features (substrate elements, temperature, fog, vapor pressure etc.) that affect these valleys, we can gain a better understanding of how they have evolved and why certain species are found there. This knowledge will provide more insights into the biogeography and development of these valley groups and help further understanding of how biotic and abiotic factors guide evolutionary processes in the *Huidobria* shrub species within the Atacama Desert.

## Acknowledgments

We thank Claudia Schütte (BIOB) for the molecular lab work as well as Prof. Dr. Heiko Schoof (INRES, Bonn) for helpful discussions and support. We appreciate the helpful comments of the reviewers. Finally, we would like to thank our colleagues in the framework of the Atacama project: Earth– Evolution at the dry Limit. This project is affiliated to the Collaborative Research Center (CRC) 1211 and was funded by the Deutsche Forschungsgemeinschaft (DFG, German Research Foundation – Projektnummer 268236062 – SFB 1211; http://sfb1211.uni-koeln.de/).

## Additional Information

### Data availability statement

All data generated or analyzed during this study are included in this published article (and its Supplementary Information files).

### Author contributions

KBF, TB, BR, DQ, TW designed the study. AS, PM conducted fieldwork. KBF, TB, DH, TW analyzed the genetic data. BR provided geological part of the paper. SF, FM analyzed climatic data. KBF wrote the manuscript with substantial support by TB, BR, TW, further contributions were made by JB, CM, MAK, DQ. All authors made edits and approved the manuscript.

## Supplementary material

**Table S1.** Rainfall difference matrix: Matrix generated from the Precipitation data in the Atacama Desert during 20 years from January 2001 to December 2021. Sum of Precipitation was calculated for each site/year, then average of all years was calculated for each site. Absolute values were used.

**Table S2.** Elevation difference matrix: Matrix generated from the elevation data of the studied area. Elevation data were extracted from ArcGIS Pro software and differences for every pairwise sites were calculated. Absolute values were used.

**Table S3.** GPS distance matrix. Matrix generated from the GPS data of the studied sites. GPS data were extracted from the KML file using ArcGIS Pro software and matrix was generated using the Geographic Distance Matrix Generator. Absolute values were used.

**Table S4.** Comparative table of the polymorphic sites in two datasets. **pop ID:** population Identifier; **N°Ind**: Number of individuals processed in the analysis; **Psites_005maf**: Polymorphic sites identified in the fileterd dataset (based on 0.05 minor allel frequency and 0.5 missing data); **Psites_orig**: Polymorphic sites identified in the firstly generated dataset (applying r = 0.65); **HN°**: Harmonic number calculated for every population; **\*Dataset.orig**: Normalized number of polymorphic sites in the original dataset, calculated by dividing the polymorphic sites number from the column “Psites_orig” by the harmonic number; **\*Dataset.005maf**: Normalized number of polymorphic sites in the filtered dataset, calculated by dividing the polymorphic sites number from the column “Psites_005maf” by the harmonic number.

**Table S5.** Table summarizing the polymorphism information in the filtered dataset (applying 0.05 minor allele frequency and 0.5 missing data parameters). **Pop ID:** Population identifier; **Private alleles**: alleles identified in only this population; **VS:** Variant sites; **PS:** Polymorphic sites; %PL: Percentage of polymorphic loci; **P:** Mean frequency of the most frequent alleles in each locus for this population.

**Table S6.** Sampling data of 186 individuals of *Huidobria chilensis* collected from 11 different locations. ID: Identifier for all collected individuals. Numbers from 800 to 810 refers to the different stations and numbers from 1 to 20 identify the individuals; S Latitude: Latitude coordinate for each sample; W Longitude: Longitude coordinate for each sample.

**Table S7.** Matrix of pairwise F_ST_ values of all population comparisons: Values calculated for the filtered SNPs data and generated by populations tool of stacks software.

**Table S8.** Evanno Table for 183 individuals of *Huidobria chilensis* (eleven populations) showing maximum delta K = 3, marked in yellow. Table was performed by Structure HARVESTER software.

**Figure S1.** Delta K plot of Evanno method test based on STRUCTURE analysis. Plot was generated by Structure HARVESTER software using all runs output of structure analysis.

**Figure S2.** STRUCTURE bar plots representing K = 3 for different simulation parameter combinations. **Admix+correlated AF**: STRUCTURE runs under admixture model and with correlated allele frequencies; **Admix+uncorrelated AF**: STRUCTURE runs under admixture model and with uncorrelated allele frequencies; **N**o **Admix+correlated AF**: STRUCTURE runs without admixture model and with correlated allele frequencies; **No Admix+uncorrelated AF**: STRUCTURE runs without admixture model and with uncorrelated allele frequencies.

**Figure S3.** Climatology of precipitation in the Atacama Desert for 1991-2020. Shown are precipitation (in mm day-1) as (a) annual mean. (b) Autumn (SON) mean. (c) Winter (DJF) mean. (d) Spring (MAM) mean. (e) Summer (JJA) mean. (f) 90th percentile in JJA. Data based on WRF high-resolution simulation driven by ERA5 re-analysis.

