## Supplementary material for "Genetic diversity of the Atacama Desert shrub *Huidobria chilensis* in the context of geography and climate"

**Table S1.** Rainfall difference matrix: Matrix generated from the Precipitation data in the Atacama Desert during 20 years from January 2001 to December 2021. Sum of Precipitation was calculated for each site/year, then average of all years was calculated for each site. Absolute values were used.

| | 800 | 801 | 802 | 803 | 804 | 805 | 806 | 807 | 808 | 809 | 810 |
|---|---|---|---|---|---|---|---|---|---|---|---|
| **800** | 0 | 0.00644 | 0.00517 | 0.00046 | 0.0087 | 0.0158 | 0.0032 | 0.0003 | 0.0012 | 0.0037 | 0.00672 |
| **801** | 0.00644 | 0 | 0.00127 | 0.00598 | 0.0023 | 0.0093 | 0.0033 | 0.0062 | 0.0076 | 0.0102 | 0.00028 |
| **802** | 0.00517 | 0.00127 | 0 | 0.00471 | 0.0035 | 0.0106 | 0.002 | 0.0049 | 0.0063 | 0.0089 | 0.00155 |
| **803** | 0.00046 | 0.00598 | 0.00471 | 0 | 0.0082 | 0.0153 | 0.0027 | 0.0002 | 0.0016 | 0.0042 | 0.00626 |
| **804** | 0.00871 | 0.00227 | 0.00354 | 0.00824 | 0 | 0.0071 | 0.0056 | 0.0084 | 0.0099 | 0.0124 | 0.00199 |
| **805** | 0.01579 | 0.00935 | 0.01062 | 0.01532 | 0.0071 | 0 | 0.0126 | 0.0155 | 0.017 | 0.0195 | 0.00907 |
| **806** | 0.00316 | 0.00329 | 0.00201 | 0.00269 | 0.0056 | 0.0126 | 0 | 0.0029 | 0.0043 | 0.0069 | 0.00356 |
| **807** | 0.00027 | 0.00617 | 0.00489 | 0.00019 | 0.0084 | 0.0155 | 0.0029 | 0 | 0.0014 | 0.004 | 0.00645 |
| **808** | 0.00116 | 0.00761 | 0.00633 | 0.00163 | 0.0099 | 0.017 | 0.0043 | 0.0014 | 0 | 0.0026 | 0.00788 |
| **809** | 0.00373 | 0.01017 | 0.0089 | 0.00419 | 0.0124 | 0.0195 | 0.0069 | 0.004 | 0.0026 | 0 | 0.01045 |
| **810** | 0.00672 | 0.00028 | 0.00155 | 0.00626 | 0.002 | 0.0091 | 0.0036 | 0.0064 | 0.0079 | 0.0104 | 0 |

**Table S2.** Elevation difference matrix: Matrix generated from the elevation data of the studied area. Elevation data were extracted from ArcGIS Pro software and differences for every pairwise sites were calculated. Absolute values were used.

| | 800 | 801 | 802 | 803 | 804 | 805 | 806 | 807 | 808 | 809 | 810 |
|---|---|---|---|---|---|---|---|---|---|---|---|
| **800** | 0 | 1018.4 | 1656.3 | 1560.8 | 604.3 | 628.7 | 302 | 334.5 | 143.4 | 12.1 | 149.8 |
| **801** | 1018.4 | 0 | 637.9 | 542.4 | 414.1 | 389.7 | 716.4 | 1352.9 | 1161.8 | 1030.5 | 868.6 |
| **802** | 1656.3 | 637.9 | 0 | 95.5 | 1052 | 1027.6 | 1354.3 | 1990.8 | 1799.7 | 1668.4 | 1506.5 |
| **803** | 1560.8 | 542.4 | 95.5 | 0 | 956.5 | 932.1 | 1258.8 | 1895.3 | 1704.2 | 1572.9 | 1411 |
| **804** | 604.3 | 414.1 | 1052 | 956.5 | 0 | 24.4 | 302.3 | 938.8 | 747.7 | 616.4 | 454.5 |
| **805** | 628.7 | 389.7 | 1027.6 | 932.1 | 24.4 | 0 | 326.7 | 963.2 | 772.1 | 640.8 | 478.9 |
| **806** | 302 | 716.4 | 1354.3 | 1258.8 | 302.3 | 326.7 | 0 | 636.5 | 445.4 | 314.1 | 152.2 |
| **807** | 334.5 | 1352.9 | 1990.8 | 1895.3 | 938.8 | 963.2 | 636.5 | 0 | 191.1 | 322.4 | 484.3 |
| **808** | 143.4 | 1161.8 | 1799.7 | 1704.2 | 747.7 | 772.1 | 445.4 | 191.1 | 0 | 131.3 | 293.2 |
| **809** | 12.1 | 1030.5 | 1668.4 | 1572.9 | 616.4 | 640.8 | 314.1 | 322.4 | 131.3 | 0 | 161.9 |
| **810** | 149.8 | 868.6 | 1506.5 | 1411 | 454.5 | 478.9 | 152.2 | 484.3 | 293.2 | 161.9 | 0 |

**Table S3.** GPS distance matrix. Matrix generated from the GPS data of the studied sites. GPS data were extracted from the KML file using ArcGIS Pro software and matrix was generated using the Geographic Distance Matrix Generator. Absolute values were used.

| | **800** | **801** | **802** | **803** | **804** | **805** | **806** | **807** | **808** | **809** | **810** |
|---|---|---|---|---|---|---|---|---|---|---|---|
| **800** | 0 | 67.26 | 82.61 | 103.49 | 106.6 | 96.69 | 91.07 | 514.3 | 192.96 | 210.62 | 124.74 |
| **801** | 67.26 | 0 | 15.75 | 50.94 | 74 | 71.15 | 85.77 | 490.71 | 182.82 | 195.36 | 119.14 |
| **802** | 82.61 | 15.75 | 0 | 41.12 | 69.6 | 69.36 | 88.23 | 482.75 | 180.03 | 191.1 | 119.38 |
| **803** | 103.49 | 50.94 | 41.12 | 0 | 34.19 | 39.74 | 65.59 | 441.63 | 142.98 | 152.25 | 88.93 |
| **804** | 106.6 | 74 | 69.6 | 34.19 | 0 | 12.1 | 37.71 | 417.79 | 110.53 | 121.68 | 54.98 |
| **805** | 96.69 | 71.15 | 69.36 | 39.74 | 12.1 | 0 | 26.99 | 423.76 | 111.67 | 124.65 | 50.58 |
| **806** | 91.07 | 85.77 | 88.23 | 65.59 | 37.71 | 26.99 | 0 | 423.33 | 103.45 | 119.98 | 35.42 |
| **807** | 514.3 | 490.71 | 482.75 | 441.63 | 417.79 | 423.76 | 423.33 | 0 | 322.68 | 303.87 | 389.88 |
| **808** | 192.96 | 182.82 | 180.03 | 142.98 | 110.53 | 111.67 | 103.45 | 322.68 | 0 | 21.52 | 68.4 |
| **809** | 210.62 | 195.36 | 191.1 | 152.25 | 121.68 | 124.65 | 119.98 | 303.87 | 21.52 | 0 | 86.01 |
| **810** | 124.74 | 119.14 | 119.38 | 88.93 | 54.98 | 50.58 | 35.42 | 389.88 | 68.4 | 86.01 | 0 |

**Table S4.** Comparative table of the polymorphic sites in two datasets. **pop ID:** population Identifier; **N°Ind**: Number of individuals processed in the analysis; **Psites_005maf**: Polymorphic sites identified in the fileterd dataset (based on 0.05 minor allel frequency and 0.5 missing data); **Psites_orig**: Polymorphic sites identified in the firstly generated dataset (applying r = 0.65); **HN°**: Harmonic number calculated for every population; ***Dataset.orig**: Normalized number of polymorphic sites in the original dataset, calculated by dividing the polymorphic sites number from the column "Psites_orig" by the harmonic number; ***Dataset.005maf**: Normalized number of polymorphic sites in the filtered dataset, calculated by dividing the polymorphic sites number from the column "Psites_005maf" by the harmonic number.

| **pop ID** | **N°Ind** | **Psites_005maf** | **Psites_orig** | **HN°** | ***Dataset.orig** | ***Dataset.005maf** |
|---|---|---|---|---|---|---|
| **805** | 20 | 15069 | 25718 | 4.253543039 | 6046 | 3543 |
| **806** | 20 | 14582 | 24117 | 4.253543039 | 5670 | 3428 |
| **809** | 20 | 13969 | 23811 | 4.253543039 | 5598 | 3284 |
| **808** | 19 | 13855 | 23107 | 4.201586224 | 5500 | 3298 |
| **804** | 17 | 13426 | 19606 | 4.088798226 | 4795 | 3284 |
| **803** | 19 | 13143 | 18718 | 4.201586224 | 4455 | 3128 |
| **810** | 4 | 8251 | 11413 | 2.592857143 | 4402 | 3182 |
| **802** | 9 | 9673 | 12971 | 3.439552523 | 3771 | 2812 |
| **801** | 15 | 9995 | 13874 | 3.961653798 | 3502 | 2523 |
| **800** | 20 | 10605 | 14874 | 4.253543039 | 3497 | 2493 |
| **807** | 20 | 7487 | 12023 | 4.253543039 | 2827 | 1760 |

**Table S5.** Table summarizing the polymorphism information in the filtered dataset (applying 0.05 minor allele frequency and 0.5 missing data parameters). **Pop ID:** Population identifier; **Private alleles**: alleles identified in only this population; **VS:** Variant sites; **PS:** Polymorphic sites; %PL: Percentage of polymorphic loci; **P:** Mean frequency of the most frequent alleles in each locus for this population.

| Pop ID | Private Alleles | VS | PS | %PL | P |
|---|---|---|---|---|---|
| **800** | 50 | 16127 | 10605 | 65.75929 | 0.85637 |
| **801** | 4 | 16564 | 9995 | 60.3417 | 0.8618 |
| **802** | 1 | 16478 | 9673 | 58.70251 | 0.86022 |
| **803** | 0 | 15957 | 13143 | 82.36511 | 0.81489 |
| **804** | 0 | 16246 | 13426 | 82.64188 | 0.81464 |
| **805** | 0 | 17120 | 15069 | 88.01986 | 0.80969 |
| **806** | 0 | 17191 | 14582 | 84.82345 | 0.81241 |
| **807** | 452 | 15876 | 7487 | 47.15923 | 0.89386 |
| **808** | 1 | 16746 | 13855 | 82.73618 | 0.818 |
| **809** | 1 | 16830 | 13969 | 83.00059 | 0.81859 |
| **810** | 0 | 14974 | 8251 | 55.10218 | 0.85163 |

**Table S6.** Sampling data of 186 individuals of *Huidobria chilensis* collected from 11 different locations. ID: Identifier for all collected individuals. Numbers from 800 to 810 refers to the different stations and numbers from 1 to 20 identify the individuals; S Latitude: Latitude coordinate for each sample; W Longitude: Longitude coordinate for each sample.

| ID | S Latitude | W Longitude |
|---|---|---|
| **800-1** | -27.36840298 | -70.435444 |
| **800-2** | -27.36837097 | -70.435444 |
| **800-3** | -27.36808104 | -70.43540703 |
| **800-4** | -27.36756999 | -70.43562303 |
| **800-5** | -27.37181702 | -70.43503999 |
| **800-6** | -27.371771 | -70.43502297 |
| **800-7** | -27.37170302 | -70.43504703 |
| **800-8** | -27.37016302 | -70.43520897 |
| **800-9** | -27.37006201 | -70.435201 |
| **800-10** | -27.36912098 | -70.43521198 |
| **800-11** | -27.37419597 | -70.43373803 |
| **800-12** | -27.37397301 | -70.434006 |
| **800-13** | -27.37438699 | -70.43340703 |
| **800-14** | -27.37445598 | -70.43323302 |
| **800-15** | -27.37446402 | -70.43316999 |
| **800-16** | -27.37498797 | -70.43169301 |

| | | |
|---|---|---|
| **800-17** | -27.37507699 | -70.43165102 |
| **800-18** | -27.37542702 | -70.43123402 |
| **800-19** | -27.37586204 | -70.43069003 |
| **800-20** | -27.37603898 | -70.43056598 |
| **801-1** | -27.13695399 | -69.807605 |
| **801-2** | -27.13696104 | -69.80763501 |
| **801-3** | -27.13727502 | -69.80750299 |
| **801-4** | -27.13750997 | -69.80788303 |
| **801-5** | -27.137501 | -69.807908 |
| **801-6** | -27.13778699 | -69.80792804 |
| **801-7** | -27.13850004 | -69.80778102 |
| **801-8** | -27.13835704 | -69.80825803 |
| **801-9** | -27.13784499 | -69.80930602 |
| **801-10** | -27.13715298 | -69.80874997 |
| **801-11** | -27.13634899 | -69.80837697 |
| **801-12** | -27.13609703 | -69.80852902 |
| **801-13** | -27.13601799 | -69.80830598 |
| **801-14** | -27.13602603 | -69.80794304 |
| **801-15** | -27.13587801 | -69.80831604 |
| **802-1** | -27.052163 | -69.68000703 |
| **802-2** | -27.052235 | -69.67993897 |
| **802-3** | -27.052465 | -69.67981299 |
| **802-4** | -27.05255996 | -69.67968098 |
| **802-5** | -27.05296003 | -69.67983797 |
| **802-6** | -27.05280304 | -69.68002698 |
| **802-7** | -27.052667 | -69.68007602 |
| **802-8** | -27.05258897 | -69.68049997 |
| **802-9** | -27.05255804 | -69.68070097 |
| **803-1** | -26.685717 | -69.72735398 |
| **803-2** | -26.67748204 | -69.73628397 |
| **803-3** | -26.67666203 | -69.73676501 |
| **803-4** | -26.67699999 | -69.73608498 |
| **803-5** | -26.67700603 | -69.73564301 |
| **803-6** | -26.67813901 | -69.73358399 |
| **803-7** | -26.67830899 | -69.733298 |
| **803-8** | -26.67884904 | -69.73234196 |
| **803-9** | -26.68481897 | -69.727228 |
| **803-10** | -26.68397902 | -69.72650497 |
| **803-11** | -26.68222896 | -69.72649198 |
| **803-12** | -26.68216601 | -69.72654202 |
| **803-13** | -26.68215796 | -69.72641998 |
| **803-14** | -26.68535398 | -69.72610398 |
| **803-15** | -26.69500901 | -69.720757 |
| **803-16** | -26.69502603 | -69.72091097 |
| **803-17** | -26.69467399 | -69.72126301 |
| **803-18** | -26.69186798 | -69.72216901 |
| **803-19** | -26.69144 | -69.72379804 |

| | | |
|---|---|---|
| **803-20** | -26.69116197 | -69.72423096 |
| **804-1** | -26.49403604 | -69.99631198 |
| **804-2** | -26.49204802 | -69.99766398 |
| **804-3** | -26.49173797 | -69.99786599 |
| **804-4** | -26.491743 | -69.99797001 |
| **804-5** | -26.49162197 | -69.99824996 |
| **804-6** | -26.49154896 | -69.99827502 |
| **804-7** | -26.49222496 | -70.00006297 |
| **804-8** | -26.49230802 | -69.99993204 |
| **804-9** | -26.49305401 | -70.00078197 |
| **804-10** | -26.49304697 | -70.00081499 |
| **804-11** | -26.49121201 | -70.00202601 |
| **804-12** | -26.49120304 | -70.00200296 |
| **804-13** | -26.49018799 | -70.00506797 |
| **804-14** | -26.49015698 | -70.00512103 |
| **804-15** | -26.49067297 | -70.00578102 |
| **804-16** | -26.49179799 | -70.00565596 |
| **804-17** | -26.49209403 | -70.00538598 |
| **805-1** | -26.55374704 | -70.09888904 |
| **805-2** | -26.55397897 | -70.09851303 |
| **805-3** | -26.554804 | -70.098292 |
| **805-4** | -26.55573498 | -70.09818303 |
| **805-5** | -26.55576699 | -70.09791397 |
| **805-6** | -26.55484901 | -70.09843801 |
| **805-7** | -26.55362299 | -70.09898099 |
| **805-8** | -26.55289602 | -70.09911904 |
| **805-9** | -26.549023 | -70.10103296 |
| **805-10** | -26.54936599 | -70.10097597 |
| **805-11** | -26.54947302 | -70.10094998 |
| **805-12** | -26.54981601 | -70.10106004 |
| **805-13** | -26.55009898 | -70.100895 |
| **805-14** | -26.55062797 | -70.10085904 |
| **805-15** | -26.55078596 | -70.10087597 |
| **805-16** | -26.53606902 | -70.11147598 |
| **805-17** | -26.53668903 | -70.11088103 |
| **805-18** | -26.50837496 | -70.153531 |
| **805-19** | -26.50340198 | -70.16213897 |
| **805-20** | -26.50422902 | -70.16074204 |
| **806-1** | -26.55271204 | -70.36976097 |
| **806-2** | -26.55181199 | -70.36619397 |
| **806-4** | -26.55191702 | -70.366415 |
| **806-5** | -26.551742 | -70.36415599 |
| **806-6** | -26.55168199 | -70.36394301 |
| **806-7** | -26.55144897 | -70.364056 |
| **806-8** | -26.55115401 | -70.36376196 |
| **806-9** | -26.55110699 | -70.36357303 |
| **806-10** | -26.550951 | -70.36303902 |

| | | |
|---|---|---|
| **806-11** | -26.55062696 | -70.36286602 |
| **806-12** | -26.55048899 | -70.36164 |
| **806-13** | -26.55038598 | -70.36136096 |
| **806-14** | -26.55029202 | -70.36111102 |
| **806-15** | -26.55021197 | -70.360274 |
| **806-16** | -26.55020904 | -70.360102 |
| **806-17** | -26.55142701 | -70.36455698 |
| **806-18** | -26.551325 | -70.36497398 |
| **806-19** | -26.55162801 | -70.36738001 |
| **806-20** | -26.55111001 | -70.36777203 |
| **807-1** | -22.75025697 | -70.28345503 |
| **807-2** | -22.74864304 | -70.283205 |
| **807-3** | -22.74863097 | -70.28292202 |
| **807-4** | -22.74778901 | -70.28343701 |
| **807-5** | -22.74746102 | -70.28303904 |
| **807-6** | -22.75032504 | -70.28282102 |
| **807-7** | -22.75023401 | -70.27709702 |
| **807-8** | -22.749859 | -70.27782097 |
| **807-9** | -22.74996202 | -70.27789297 |
| **807-10** | -22.75012597 | -70.27795499 |
| **807-11** | -22.75023703 | -70.27799598 |
| **807-12** | -22.75056903 | -70.27811802 |
| **807-13** | -22.75101998 | -70.27830603 |
| **807-14** | -22.75070298 | -70.27901002 |
| **807-15** | -22.75068998 | -70.27907096 |
| **807-16** | -22.75062603 | -70.27930004 |
| **807-17** | -22.75058596 | -70.27936298 |
| **807-18** | -22.750563 | -70.27942501 |
| **807-19** | -22.75059703 | -70.27952501 |
| **807-20** | -22.75036301 | -70.27954697 |
| **808-1** | -25.63783402 | -70.55606503 |
| **808-2** | -25.64399103 | -70.56681398 |
| **808-3** | -25.65045699 | -70.62969703 |
| **808-4** | -25.65021903 | -70.63203097 |
| **808-5** | -25.65125202 | -70.621897 |
| **808-6** | -25.65164102 | -70.620718 |
| **808-7** | -25.64917297 | -70.59811 |
| **808-8** | -25.64920901 | -70.595737 |
| **808-9** | -25.64882001 | -70.59426597 |
| **808-10** | -25.64896996 | -70.59383296 |
| **808-11** | -25.64889402 | -70.59336802 |
| **808-12** | -25.64834501 | -70.59071699 |
| **808-13** | -25.64504597 | -70.57775304 |
| **808-14** | -25.64517204 | -70.57678401 |
| **808-15** | -25.64494799 | -70.57335497 |
| **808-16** | -25.64484296 | -70.57326796 |
| **808-17** | -25.64475001 | -70.57285398 |

| | | |
|---|---|---|
| **808-18** | -25.64480198 | -70.57269104 |
| **808-19** | -25.64353203 | -70.56643596 |
| **808-20** | -25.626887 | -70.54647496 |
| **809-1** | -25.47626998 | -70.43972497 |
| **809-2** | -25.47798098 | -70.440493 |
| **809-3** | -25.48101498 | -70.43874596 |
| **809-4** | -25.48174697 | -70.43825101 |
| **809-5** | -25.48331799 | -70.43683204 |
| **809-6** | -25.48342704 | -70.436596 |
| **809-7** | -25.48369702 | -70.43667102 |
| **809-8** | -25.48395703 | -70.43669399 |
| **809-9** | -25.48481299 | -70.43588698 |
| **809-10** | -25.48478197 | -70.43599896 |
| **809-11** | -25.48489697 | -70.43515901 |
| **809-12** | -25.484642 | -70.434654 |
| **809-13** | -25.48456103 | -70.43446097 |
| **809-14** | -25.48444502 | -70.43368799 |
| **809-15** | -25.48447603 | -70.43367399 |
| **809-16** | -25.48454703 | -70.43360299 |
| **809-17** | -25.48483201 | -70.43231796 |
| **809-18** | -25.48477401 | -70.43198302 |
| **809-19** | -25.48488004 | -70.43152303 |
| **809-20** | -25.48492597 | -70.43112002 |
| **810-1** | -26.24811301 | -70.47435899 |
| **810-2** | -26.24805702 | -70.47430803 |
| **810-3** | -26.24807303 | -70.47426403 |
| **810-4** | -26.24811703 | -70.47339499 |
| **810-5** | -26.24811301 | -70.47336398 |

**Table S7.** Matrix of pairwise $F_{ST}$ values of all population comparisons: Values calculated for the filtered SNPs data and generated by populations tool of stacks software.

| | **800** | **801** | **802** | **803** | **804** | **805** | **806** | **807** | **808** | **809** | **810** |
|---|---|---|---|---|---|---|---|---|---|---|---|
| **800** | 0.000000 | 0.121919 | 0.1142370 | 0.1144520 | 0.1171970 | 0.1037440 | 0.1084960 | 0.251909 | 0.1271200 | 0.1274990 | 0.1279860 |
| **801** | 0.121919 | 0.000000 | 0.0835200 | 0.1128460 | 0.1174780 | 0.1026090 | 0.1091910 | 0.268930 | 0.1300410 | 0.1304440 | 0.1509320 |
| **802** | 0.114237 | 0.083520 | 0.0000000 | 0.0976324 | 0.1027970 | 0.0865424 | 0.0929452 | 0.261351 | 0.1156840 | 0.1156330 | 0.1613780 |
| **803** | 0.114452 | 0.112846 | 0.0976324 | 0.0000000 | 0.0325957 | 0.0315680 | 0.0415024 | 0.192316 | 0.0724250 | 0.0730942 | 0.0600449 |
| **804** | 0.117197 | 0.117478 | 0.1027970 | 0.0325957 | 0.0000000 | 0.0287403 | 0.0389912 | 0.194582 | 0.0693210 | 0.0701497 | 0.0597441 |
| **805** | 0.103744 | 0.102609 | 0.0865424 | 0.0315680 | 0.0287403 | 0.0000000 | 0.0262937 | 0.176745 | 0.0569646 | 0.0580483 | 0.0441197 |
| **806** | 0.108496 | 0.109191 | 0.0929452 | 0.0415024 | 0.0389912 | 0.0262937 | 0.0000000 | 0.182840 | 0.0626534 | 0.0636049 | 0.0493377 |
| **807** | 0.251909 | 0.268930 | 0.2613510 | 0.1923160 | 0.1945820 | 0.1767450 | 0.1828400 | 0.000000 | 0.1809550 | 0.1780470 | 0.2305950 |
| **808** | 0.127120 | 0.130041 | 0.1156840 | 0.0724250 | 0.0693210 | 0.0569646 | 0.0626534 | 0.180955 | 0.0000000 | 0.0275075 | 0.0684451 |
| **809** | 0.127499 | 0.130444 | 0.1156330 | 0.0730942 | 0.0701497 | 0.0580483 | 0.0636049 | 0.178047 | 0.0275075 | 0.0000000 | 0.0677662 |
| **810** | 0.127986 | 0.150932 | 0.1613780 | 0.0600449 | 0.0597441 | 0.0441197 | 0.0493377 | 0.230595 | 0.0684451 | 0.0677662 | 0.0000000 |

**Table S8.** Evanno Table for 183 individuals of *Huidobria chilensis* (eleven populations) showing maximum delta K = 3, marked in yellow. Table was performed by Structure HARVESTER software.

| **K** | **Reps** | **Mean LnP(K)** | **Stdev LnP(K)** | **Ln'(K)** | **\|Ln''(K)\|** | **Delta K** |
|---|---|---|---|---|---|---|
| **2** | 10 | -2966509.57 | 18.9272 | NA | NA | NA |
| **3** | 10 | -2797018.36 | 28.8639 | 169491.2 | 78345.71 | 2714.311 |
| **4** | 10 | -2705872.86 | 3659.352 | 91145.5 | 40108.11 | 10.96044 |
| **5** | 10 | -2654835.47 | 66.3459 | 51037.39 | 27164.46 | 409.437 |
| **6** | 10 | -2576633.62 | 165.654 | 78201.85 | NA | NA |

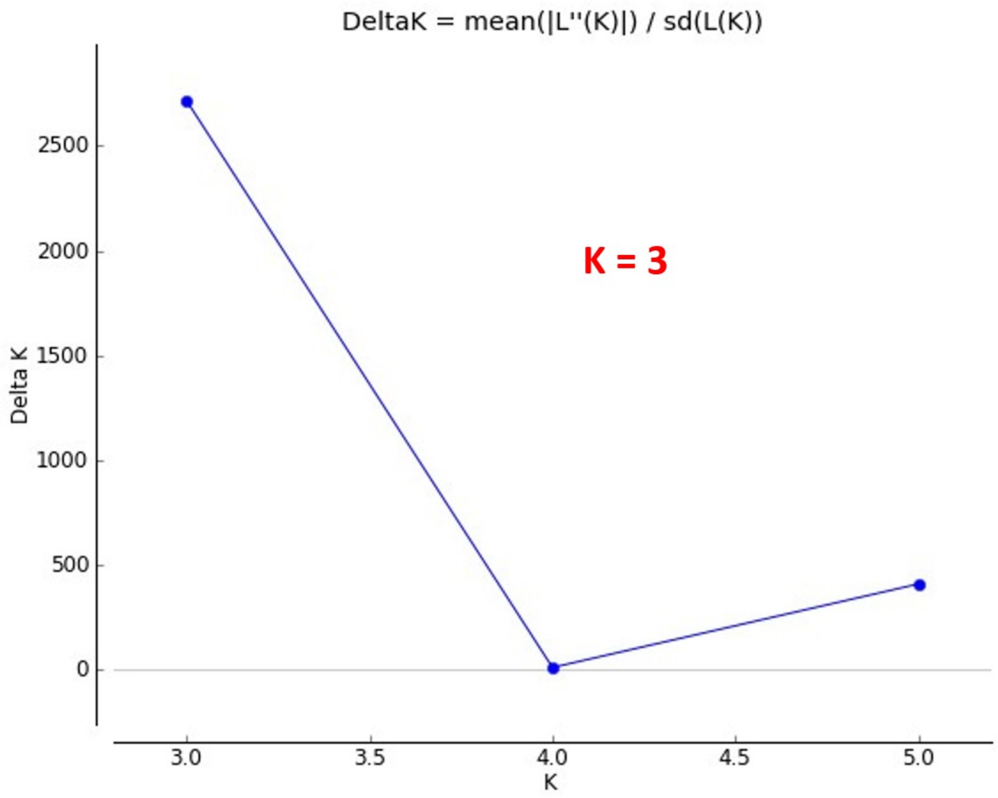


**Figure S1.** Delta K plot of Evanno method test based on STRUCTURE analysis. Plot was generated by Structure HARVESTER software using all runs output of structure analysis.

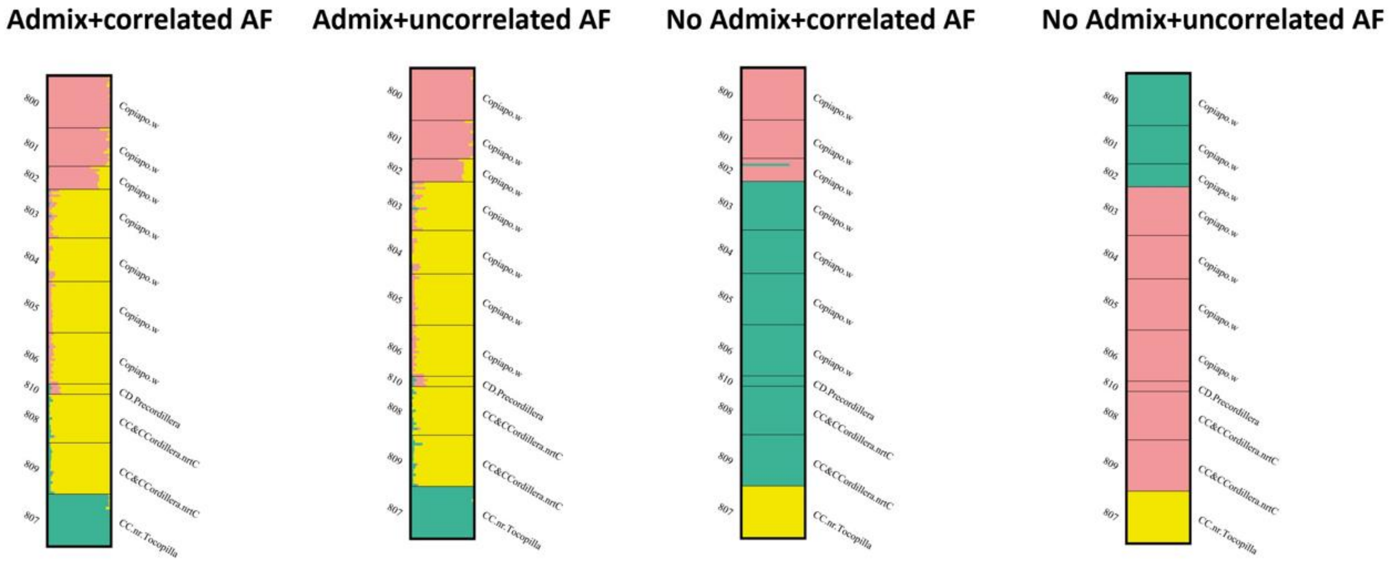


**Figure S2.** STRUCTURE bar plots representing K = 3 for different simulation parameter combinations. **Admix+correlated AF**: STRUCTURE runs under admixture model and with correlated allele frequencies; **Admix+uncorrelated AF**: STRUCTURE runs under admixture model and with uncorrelated allele frequencies; **N**o **Admix+correlated AF**: STRUCTURE runs without admixture model and with correlated allele frequencies; **No Admix+uncorrelated AF**: STRUCTURE runs without admixture model and with uncorrelated allele frequencies.

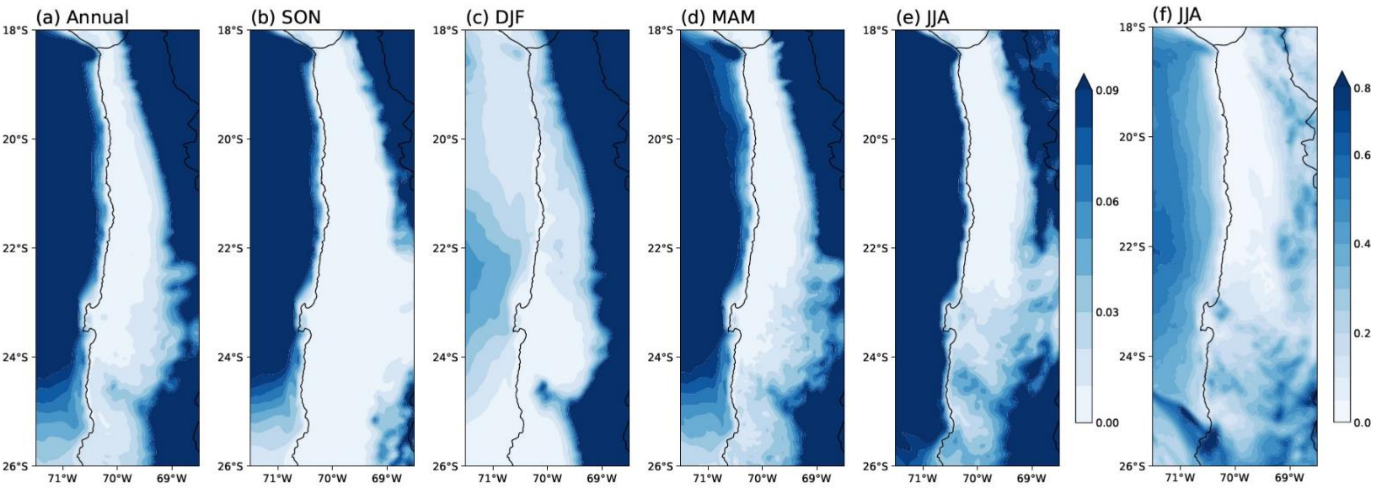


**Figure S3.** Climatology of precipitation in the Atacama Desert for 1991-2020. Shown are precipitation (in mm day-1) as (a) annual mean. (b) Autumn (SON) mean. (c) Winter (DJF) mean. (d) Spring (MAM) mean. (e) Summer (JJA) mean. (f) 90th percentile in JJA. Data based on WRF high-resolution simulation driven by ERA5 re-analysis.
